# Enhanced PI3K/Akt Signaling in Response to p85α Loss is Regulated by Endosomal PI3Kα and PI3P

**DOI:** 10.1101/2024.01.10.575035

**Authors:** Narendra Thapa, Mo Chen, Vincent L. Cryns, Richard Anderson

**Affiliations:** School of Medicine and Public Health, University of Wisconsin-Madison; 1111 Highland Ave, Madison, WI 53705, USA; Department of Medicine, School of Medicine and Public Health, University of Wisconsin-Madison; 1111 Highland Avenue, Madison, WI 53705, USA

**Keywords:** PI3K, p85α, p110α, endosomes, Akt, MAP4, PI3P

## Abstract

PI3Kα is a heterodimer of p110α catalytic subunit and p85 adaptor subunit that is activated by agonist-stimulated receptor tyrosine kinases. Although the interaction of p85α with activated receptors recruits p110α to membranes, studies have demonstrated that p85α loss, which occurs commonly in cancer, paradoxically promotes agonist-stimulated PI3K/Akt signaling. We recently demonstrated that p110α localizes to microtubules via MAP4, facilitating its interaction with activated receptor kinases in endosomes to initiate PI3K/Akt signaling. Here, we demonstrate that in response to agonist stimulation and p85α knock down the residual p110α, coupled predominantly to p85β, exhibits enhanced microtubule localization, MAP4 binding and interaction with endosomal receptor tyrosine kinases, thereby augmenting PI3K/Akt signaling. The interaction of the C2 domain of p110α with PI3P is required for recruiting p110α into endomembranes and enhancing PI3K/Akt signaling. These findings provide a mechanism for the augmented agonist-stimulated PI3K/Akt signaling upon p85α loss and point to novel therapeutic targets for cancer.

**IN BRIEF:** This study provides the comprehensive mechanism for p85α loss induced and receptor tyrosine kinase stimulated PI3K/Akt signaling.

## INTRODUCTION

The class IA phosphatidylinositol (PI) 3-kinase (PI3K) responsible for activating PI3K/Akt signaling downstream of agonist-stimulated receptor tyrosine kinases is a heterodimer composed of adaptor (p85α, p55α, p50α, p85β and p55γ) and catalytic (p110α, p110β and p110δ) subunits^1, 2^. Src homology 2 (SH2) domain-mediated recruitment of the p85 adaptor subunit to phosphorylated YXXM motifs on activated receptor tyrosine kinases liberates the intermolecular constraints imposed on the p110 catalytic subunit allowing it to the interact with the membrane surface to catalyze PI4,5P_2_ phosphorylation generating PI3,4,5P_3_^3^. PI3,4,5P_3_, in turn, recruits Akt and the Akt-activating kinases PDK1 and mTORC2 (Sin1subunit) via their pleckstrin homology domains to activate Akt by catalyzing Akt phosphorylation on T^308^ and S^473^, respectively^4^.

Despite a role for p85 in recruiting/activating the p110α catalytic subunit, heterozygous deletion of the p85α gene in mice paradoxically upregulates PI3K/Akt signaling downstream of the insulin receptor^5–7^. Similarly, liver-specific deletion of the p85α gene leads to increased PI3K/Akt signaling and hepatocellular carcinoma^8^. Moreover, p85α mRNA and protein levels are often downregulated in breast cancer tissues, implicating p85α as a tumor suppressor^8, 9^. Several studies have demonstrated decreased p85α expression but increased p85β expression in cancer, suggesting opposing functions for the p85 isoforms^10–12^. Although several potential explanations for the paradoxical increase in agonist-stimulated PI3K/Akt signaling in response to p85α loss^1, 9, 12^ have been postulated, the underlying mechanisms remain largely unknown. Given the frequent loss of p85α expression in cancer and its tumor suppressive function^8, 9^, the elucidation of these mechanisms could lead to new therapeutic targets for cancer.^8^

In PI3Kα heterodimers, the membrane recruitment of the p110α catalytic subunit is critical as induced membrane-targeting of p110α via myristylation drives constitutive PI3K/Akt signaling and is highly oncogenic^13^. Although the p85 adaptor subunits recruit p110α to activated receptor tyrosine kinases at the plasma membrane or endomembrane, multiple domains of p110α, including the Ras-binding domain (RBD), the C2 domain and the catalytic domain can directly or indirectly recruit p110 to the membrane^3, 14, 15^. Specifically, the C2 domain is a well-defined membrane-binding module present in the catalytic subunit of all PI3Ks (class I, II and III) and many other proteins such as PTEN and PKC ^3, 16, 17^. However, the precise role of the C2 domain in of PI3Kα membrane-targeting in response to agonist-stimulated PI3K/Akt signaling remains poorly understood.

Previously, we introduced the concept of IQGAP1 and MAP4 scaffolding of PI3Kα in the regulation of agonist-stimulated PI3K/Akt signaling pathway ^18, 19^. Specifically, PI3Kα is recruited by a MAP4-dependent mechanism to the endosomal compartment where it binds activated receptor tyrosine kinases to initiate PI3K-Akt signaling^19^ and IQGAP1 is a component of this complex ^18, 20^. In the present study, we discovered that p85α knockdown (KD) augments agonist-stimulated PI3K/Akt signaling predominantly at endosomal membranes. Although the expression level of p110α is substantially reduced upon p85α KD, the residual p110α, coupled largely to p85β, localizes along microtubules by binding MAP4, resulting in its recruitment to endosomal membranes and association with activated EGFR. Several basic amino acid residues in the C2 domain of p110α mediate its interaction with the endosomal phosphoinositide phosphatidylinositol 3-phosphate (PI3P), a key step in recruiting p110α to endomembranes and activating PI3K/Akt signaling. Overall, these results provide a comprehensive mechanism for the seemingly paradoxical enhanced activation of agonist-stimulated PI3K/Akt signaling in the setting of p85α loss.

## RESULTS

### p85α Adaptor Subunit KD Promotes Akt Activation Downstream of Multiple Receptor Tyrosine Kinases

We first examined the effect of p85α KD on agonist-stimulated Akt activation using siRNAs that target all the transcriptional variants of p85α (p85α, p55α, and p50α). KD of p85α enhanced Akt activation by multiple agonists as determined by western blot (WB) of pAktS^473^ levels (**Fig. 1A, B, C and D**). Moreover, three individual siRNAs targeting p85α similarly promoted EGF-stimulated Akt activation (**Fig. 1E)**. p85α KD promoted EGF-stimulated Akt activation in multiple cell types (**Suppl. Fig. S1A-C**). Collectively, these results indicate that p85α KD augments PI3K/Akt signaling downstream of multiple receptor tyrosine kinases ^5, 6^.

**Figure 1:**
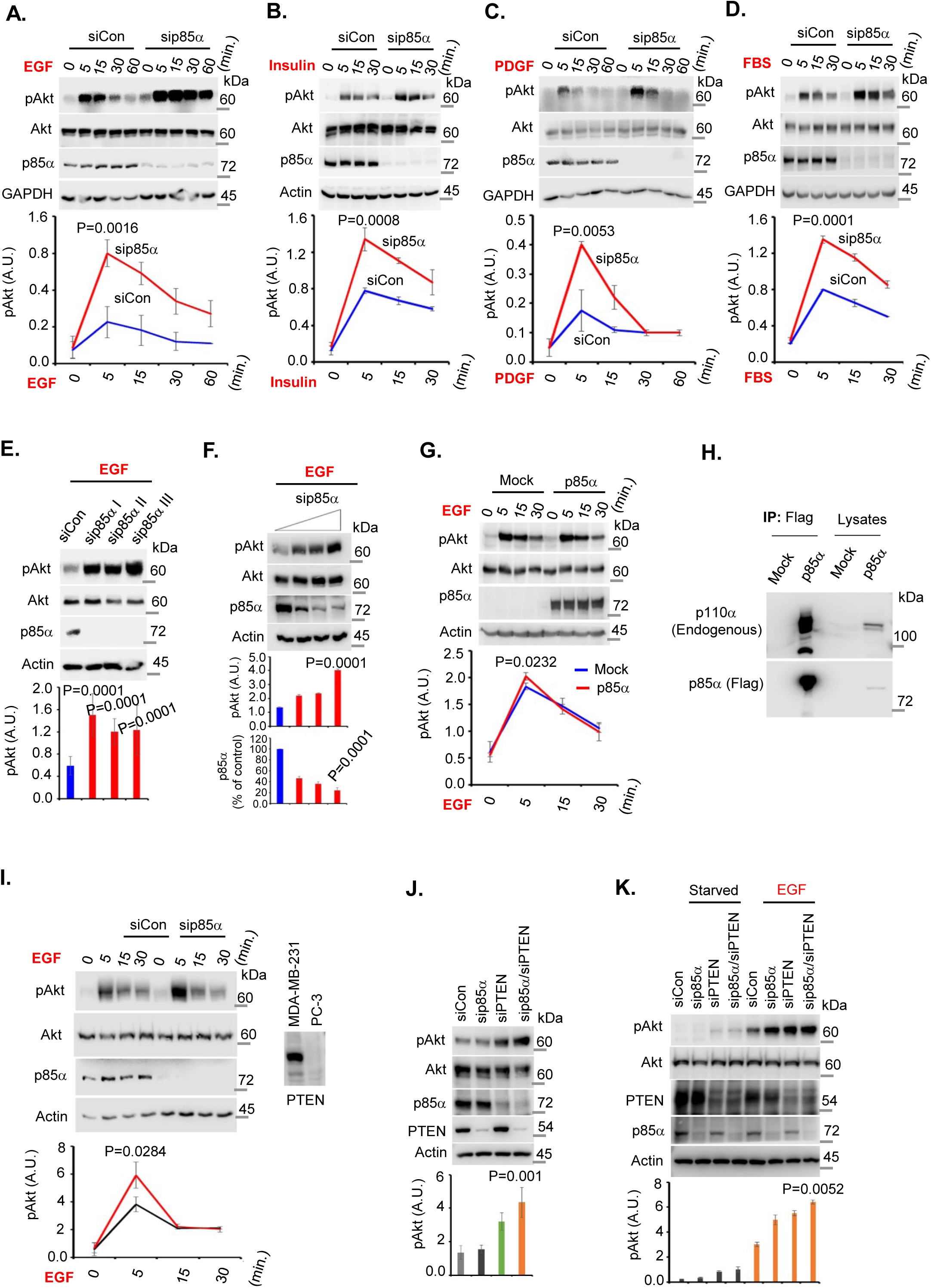
p85α Adaptor Subunit Knock Down Promotes Akt Activation Downstream of Activated Receptor Tyrosine Kinases. **A, B, C, D, KD of p85α promotes agonist-stimulated Akt activation.** MDA-MB-231 cells were transfected with either control siRNAs or siRNAs targeting all the transcriptional variants of p85α. 48-72 hours post-transfection, cells were stimulated with EGF, insulin, PDGF or FBS for different time intervals before harvesting the cells. phospho-Akt and p85α levels were analyzed by Western blot (WB) using antibodies specific for phospho-AktS^473^ and p85α. The data represent the mean ± SD from three independent experiments. The indicated p-value is for the 5-minute time point for p85α siRNA-versus control siRNA-treated cells. **E, three individual siRNAs targeting p85α show similar effect in inducing EGF-stimulated Akt activation.** MDA-MB-231 cells were transfected with control siRNAs or one of three individual siRNAs targeting endogenous p85α. 48-72 hours post-transfection, cells were stimulated with EGF for 5 minutes before harvesting the cells. Phospho-AktS^473^ and p85α levels were analyzed by WB. The data represent the mean ± SD from three independent experiments. The indicated p-value is for control individual p85α siRNA-versus control siRNA-treated cells. **F, the extent of p85α KD correlates inversely with increased EGF-stimulated Akt activation**. MDA-MB-231 cells were transfected with control siRNAs or different amount of siRNAs targeting endogenous p85α. 48-72 hours post-transfection, cells were stimulated with EGF for 5 minutes before harvesting the cells. phospho-Akt and p85α levels were analyzed by WB. The data represent the mean ± SD from three independent experiments. **G, ectopic expression of p85α does not impair EGF-stimulated Akt activation**. Cos-7 cells were transiently transfected with empty plasmid or plasmid containing p85α (Flag-tagged p85α). 48-72 hours post-transfection, cells were stimulated with EGF for different time intervals before harvesting the cells. phospho-Akt and p85α levels were analyzed by WB. The data represents the mean ± SD from three-independent experiments. The indicated p-value is for the 5-minute time point for p85α-versus mock-transfected cells. **H, ectopically expressed p85α associates with the endogenous p110α catalytic subunit.** Cos-7 cells were transfected with empty plasmid or plasmid containing p85α (Flag-tagged p85α). 48 hours post-transfection, cells were harvested for immunoprecipitation of ectopically expressed p85α using anti-Flag antibody agarose beads. Endogenous p110α that co-immunoprecipitated (co-IPed) with ectopically expressed p85α was examined by WB using a p110α-specific antibody. **I, p85α KD increases EGF-stimulated Akt activation in PTEN-null cells.** PC-3 cells were transfected with control siRNAs or siRNAs targeting p85α. 48-72 hours post-transfection, cells were stimulated with EGF before harvesting to examine the phospho-Akt and p85α levels by WB. The data represent the mean ± SD from three independent experiments. The indicated p-value is for the 5-minute time point for p85α siRNA-versus control siRNA-treated cells. **J, K, simultaneous KD of p85α and PTEN enhances EGF-stimulated Akt activation.** MDA-MB-231 cells were transfected with control siRNAs or siRNAs targeting endogenous p85α and/or PTEN. 48-72 hours post-transfection, cells were either unstimulated (**J**) or stimulated with EGF (**K**) for 5 minutes before cell harvesting. phospho-Akt and p85α levels were analyzed by WB using antibodies specific for phospho-Akt and p85α. The data represent the mean ± SD from three independent experiments. The indicated p-value is for p85α KD versus combined p85α and PTEN KD.

The p85 adaptor and p110 catalytic subunit are present in 1:1 stoichiometry in the PI3K enzyme^21, 22^. However, some studies indicate that the p85 adaptor subunit is more abundant than the p110 catalytic subunit (10-30% more depending upon cell types and tissues). Moreover, the free p85α adaptor subunit has been postulated to compete with PI3Kα heterodimers for binding to activated receptor tyrosine kinases, resulting in impaired PI3K/Akt signaling^1, 22^. To investigate whether this mechanism contributes to enhanced Akt activation in response to p85α KD, we first performed a dose-response experiment using increasing concentrations of p85α siRNAs. The degree of p85α KD correlated inversely with EGF-stimulated pAKT levels, with the most robust activation noted when p85α KD was > 90% **(Fig. 1F)**. We next examined the effect of ectopically overexpressing p85α on EGF-stimulated PI3K/Akt signaling. Robust overexpression of p85α had no effect on EGF-stimulated Akt activation (**Fig. 1G**) despite efficient co-immunoprecipitation of the endogenous p110α subunit with Flag-tagged p85α (**Fig. 1H**). These results argue against the concept that excess p85α adaptor subunits compete with the PI3Kα heterodimers for activated receptor tyrosine kinases in the cellular models we examined.

p85α binds PTEN and increased PI3K/Akt signaling upon p85α loss has been attributed to compromised PTEN recruitment to membranes to antagonize PI3K/Akt signaling^23^. To investigate this possibility, we examined the effects of p85α KD on EGF-stimulated Akt activation in PTEN-null PC-3 prostate cancer cells. p85α KD enhanced EGF-stimulated Akt activation in these PTEN-null cells (**Fig. 1H**). Furthermore, combined KD of p85α and PTEN resulted in more robust Akt activation under basal and EGF-stimulated conditions than individual KD of p85α or PTEN (**Fig. 1J, K**). Taken together, these findings demonstrate that the enhanced agonist-stimulated activation of Akt in response to p85α loss is PTEN-independent in these cellular models.

### PI3,4,5P_3_ Generation and Akt Activation induced by Agonist Stimulation and p85α KD Occur at the Endomembrane

Although the prevailing dogma is that agonist-stimulated PI3,4,5P_3_ generation and Akt activation occur exclusively at the plasma membrane^3, 24^, we recently demonstrated PI3,4,5P_3_ generation and Akt activation downstream of activated receptor tyrosine kinases occur predominantly in internal membrane compartments^19^. In response to agonist stimulation, PI3Kα distributes along microtubules, and receptor tyrosine kinases are rapid internalized and activated on endosomes to initiate agonist-stimulated PI3,4,5P_3_ generation and Akt activation in the endomembranes^19, 25^. Congruent with our WB results (**Fig. 1**), EGF stimulation and p85α KD each augmented pAkt and PI3,4,5P_3_ levels as determined by immunofluorescence (IF) (**Fig. 2A, B**). As previously^19^, we isolated plasma membrane- and endosomal membrane-enriched fractions and observed that p85α KD and EGF stimulation each activated Akt predominantly in the early endosome antigen 1 (EEA1)-enriched endosomal fraction (**Fig. 2C**). Consistent with this result, the number of EEA1- and transferrin receptor (TFR)-positive endosomes that co-localized with activated Akt and PI3,4,5P_3_ were significantly greater in cells treated with EGF or p85α siRNAs (**Fig. 2D-G; Suppl. Fig. 2A-D**). These data indicate that the increased agonist-stimulated PI3,4,5P_3_ and pAkt levels in response to p85α KD localize largely to the endosomal membrane, aligning with our previous findings^19^.

**Figure 2:**
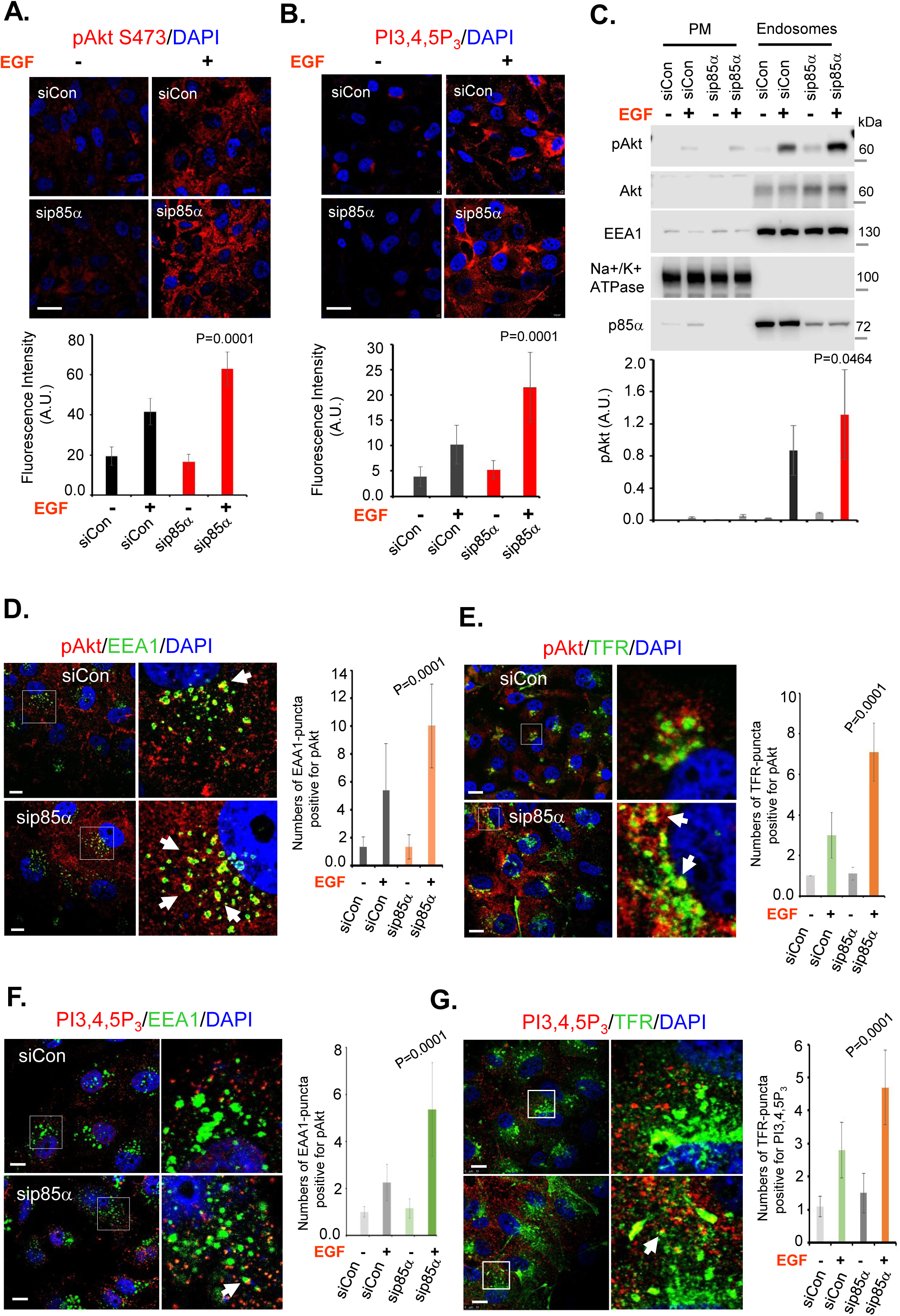
Agonist-Stimulated PI3K/Akt Signaling in Response to p85α KD Occurs at Endosomal Membranes. **A, B, IF showing activated Akt and PI3,4,5P_3_ generated upon EGF-stimulation.** MDA-MB-231 cells were transfected with control siRNAs or p85α siRNAs. 48-72 hours post-transfection, cells were stimulated with EGF for 5 minutes before fixation with 4% PFA for IF study using antibodies specific for phospho-Akt (red) and PI3,4,5P_3_ (red). IF signals were quantified in at least 20-30 cells. The p-value indicates p85α KD compared to control siRNAs upon EGF stimulation. **C, subcellular fractionation shows enrichment of activated Akt at endosomal fractions in EGF-stimulated cells.** MDA-MB-231 cells were transfected with either control siRNAs or siRNAs targeting p85α. 48-72 hours post-transfection, cells were stimulated with EGF for 5 minutes before harvesting the cells for subcellular fractionation. Phospho-Akt levels were analyzed in the plasma membrane vs endosomal fractions by WB. The data represent the mean ± SD from three independent experiments. The p-value indicates p85α KD compared with control siRNA upon EGF stimulation. **D, E, activated Akt co-localizes with endosomal markers (EEA1 and TFR).** MDA-MB-231 cells were transfected with control siRNAs or siRNAs targeting p85α. 48-72 hours post-transfection, cells were stimulated with EGF for 5 minutes before cell fixation with 4% PFA for IF using antibodies specific for phospho-Akt (red) and endosomal markers (EEA1 and TFR) (green). The number of EEA1- and TFR-uncta positive for Akt were counted in at least 20-30 cells. The p-value indicates p85α KD compared with control siRNA upon EGF stimulation. **F, G, PI3,4,5P_3_ co-localizes with endosomal markers (EEA1 and TFR).** MDA-MB-231 cells were transfected with control siRNAs or siRNAs targeting p85α. 48-72 hours post-transfection, cells were stimulated with EGF for 5 minutes before fixation with 4% PFA. The co-localization of PI3,4,5P_3_ generated with endosomal markers (EEA1 and TFR) (green) wase examined by IF using an antibody specific for PI3,4,5P_3_ (red). The number of EEA1- and TFR-puncta positive for PI3,4,5P_3_ were quantified in at least 20-30 cells. The p-value indicates p85α KD compared with control siRNA upon EGF stimulation.

### Residual p110α Catalytic Subunit is Responsible for Increased Agonist-stimulated Akt Activation in Response to p85α KD

Among class IA PI3Ks, PI3Kα and PI3Kβ are ubiquitously expressed enzymes in nonhematopoietic cells and PI3Kδ is expressed in hematopoietic cells^2^. As the p85 adaptor subunit stabilizes p110 catalytic subunits, we examined the expression level of p110α and p110β catalytic subunits upon p85α or p85β KD. p85α KD (more than 90%) resulted in ⁓60% decrease in p110α levels but only ⁓25% reduction in p110β levels (**Fig. 3A**). p85β KD resulted in ⁓20% decrease in p110α levels and ⁓10% reduction in p110β levels. To examine the contribution of p110α and p110β catalytic subunits to Akt activation in response to p85α KD and agonist stimulation, we used siRNAs to KD p85α and p110α or p110β. KD of p110α but not p110β inhibited EGF-stimulated Akt activation upon EGF stimulation (**Fig. 3B)**, indicating that the residual p110α catalytic subunit is responsible for activating PI3K/Akt signaling in response to p85α KD consistent with PI3Kα inhibitor blockade PI3K/Akt signaling p85α knocked down cells^9^. Furthermore, overexpression of p110α augments EGF-stimulated Akt activation in response to p85α KD (**Suppl. Fig. 1D**), providing additional evidence for the functional role of p110α in these events.

**Figure 3.**
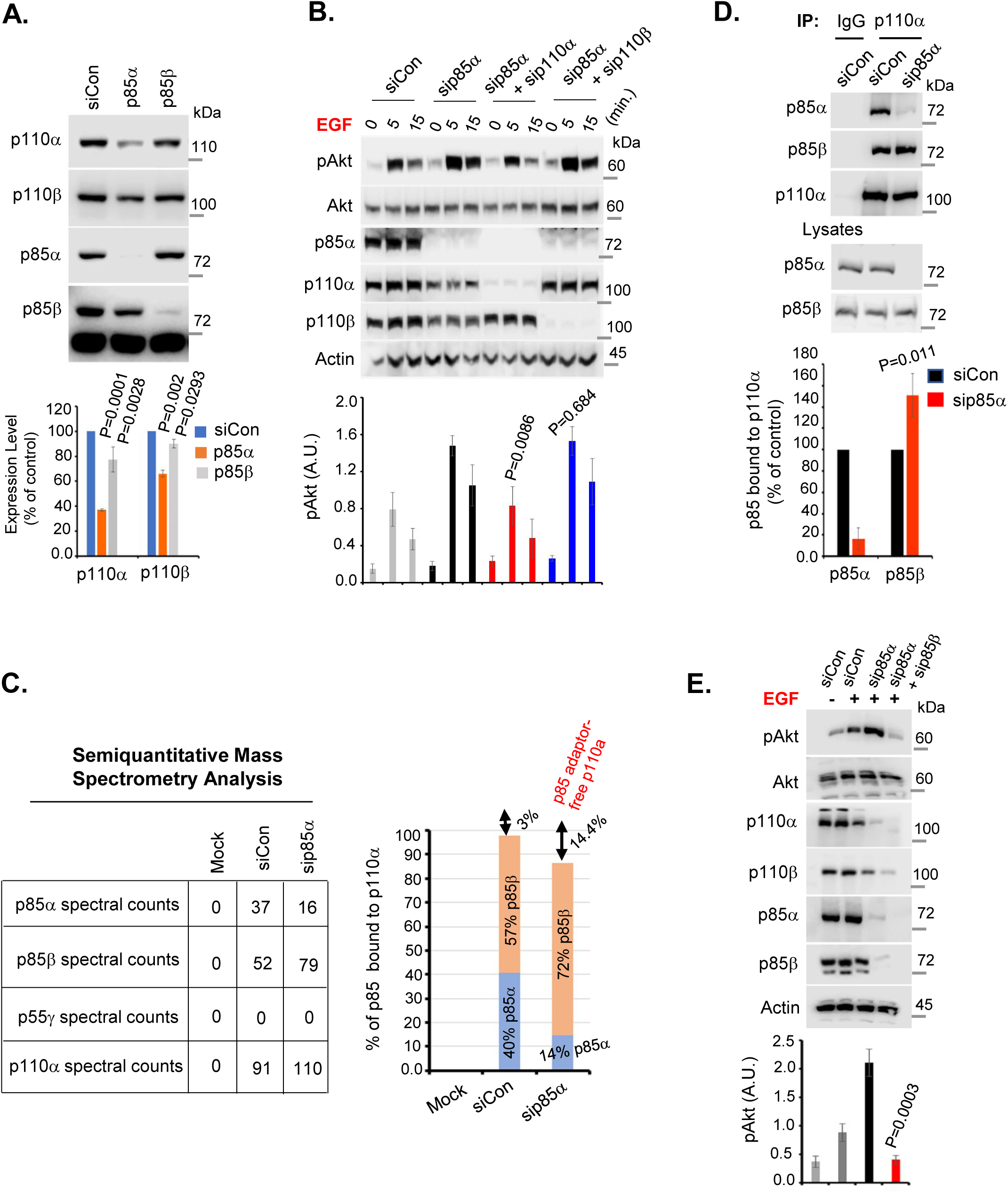
Residual p110α Catalytic Subunit is Responsible for Increased Agonist-stimulated Akt Activation Upon p85α KD. **A, p85α KD decreases p110α catalytic subunit levels.** MDA-MB-231 cells were transfected with control siRNAs or siRNAs targeting p85α or p85β. 48-72 hours post-transfection, cells were harvested and the expression level of p85α, p85β, p110α and p110β were examined by WB. The data represent the mean ± SD from three independent experiments. **B, residual p110α catalytic subunit is responsible for EGF-stimulated Akt activation upon p85α KD.** MDA-MB-231 cells were transfected with control siRNAs or siRNAs targeting p85α and p110α or p110β. 48-72 hours post-transfection, cells were stimulated with EGF for different time interval before harvesting the cells. phospho-Akt, p85α, p110α and p110β levels were analyzed by WB. The data represent the mean ± SD from three independent experiments. The significance is indicated by p-values (sip85α vs sip85α/sip110α; sip85α vs sip85α/sip110β). **C, mass spectrometry analysis of p110α-associated adaptor proteins.** MDA-MB-231 cells stably expressing HA-p110α were transfected with control siRNAs or siRNAs targeting p85α. 48-72 hours post-transfection, cells were stimulated with EGF for 5 minutes before harvesting the cells. p110α was IPed using anti-HA agarose beads and co-IPed proteins analyzed by mass spectrometry. The data table shows the spectral counts of IPed p110α catalytic subunit and its associated adaptor subunits and represents three independent mass spectrometry analyses. **D, increased association of p85β adaptor protein with p110α upon p85α KD.** MDA-MB-231 cells were transfected with control siRNAs or p85α siRNAs. 48-72 hours post-transfection, p110α was IPed and co-IPed p85 adaptor proteins were examined by WB. The data represent the mean ± SD from three independent experiments. **E, combined KD of p85α and p85β adaptor subunit abolishes EGF-stimulated Akt activation.** MDA-MB-231 cells were transfected with control siRNAs or siRNAs targeting p85α and/or p85β. 48-72 hours post-transfection, cells were stimulated with EGF for 5 minutes before harvesting the cells. phospho-Akt p85α, p85β, p110α and p110β levels were analyzed by WB. The data represent the mean ± SD from three independent experiments. The statistical significance is indicated by p-value (p85α siRNA vs sip85α/sip85β).

Next, we undertook a semiquantitative mass spectrometry approach to analyze the p110α catalytic subunit-associated adaptor subunits in response to p85α KD. The analysis of spectral counts of proteins that co-immunoprecipitated (co-IPed) with p110α indicated that p85α and p85β constituted ⁓40% and ⁓57% of p85 adaptor subunits, respectively (together ⁓97%) that associated with p110α in cells treated with non-silencing control (siCon) siRNAs (**Fig. 3C**). In response to p85α KD, p85β association with p110α increased from ⁓57% to ⁓72% with ⁓14% of p110α not associated with any p85 adaptor subunits (**Fig. 3C**). We did not detect p55γ or its association with p110α in the cells examined. These results were validated by co-IP experiments which revealed increased p85β association with residual p110α in response to p85α KD (**Fig. 3D**). However, KD of both p85α and p85β adaptor proteins almost completely abolished p110α expression and EGF-stimulated Akt activation (**Fig. 3E**). These results indicate that p110α retains competency for receptor tyrosine kinases in response to p85α KD likely through increased coupling with p85β.

### Residual p110α Localizes to Microtubules, Integrates into Endosomes, and Associates with Receptor Kinases in Response to p85α KD

We recently demonstrated PI3Kα localizes to microtubules via a direct interaction of p110α with the microtubule-associated protein 4 (MAP4)^19^. MAP4 serve as a scaffolding molecule to recruit PI3Kα vesicles along microtubules, facilitating its incorporation into endosomes containing active receptors^19^. Consistent with these results, residual p110α localized along microtubules following p85α KD **(Suppl. Fig. 3A**) and co-localized with MAP4 in a pattern indistinguishable from cells treated with siCon (**Fig. 4A**, upper panel). Furthermore, p85α KD enhanced the interaction of residual p110α with MAP4 (**Fig. 4A**, lower panel; **Suppl. Fig. 3B, C**). Following p85α KD, the residual p110α exhibited enhanced co-localization with the endosomal markers EEA1 and TFR in response to EGF stimulation (**Fig. 4B, C; Suppl. Fig. 4A, B, C**). These results indicate that p85α KD increases agonist-stimulated p110α localization along microtubules and MAP4 binding, resulting in its enhanced incorporation in endosomal membranes.

**Figure 4.**
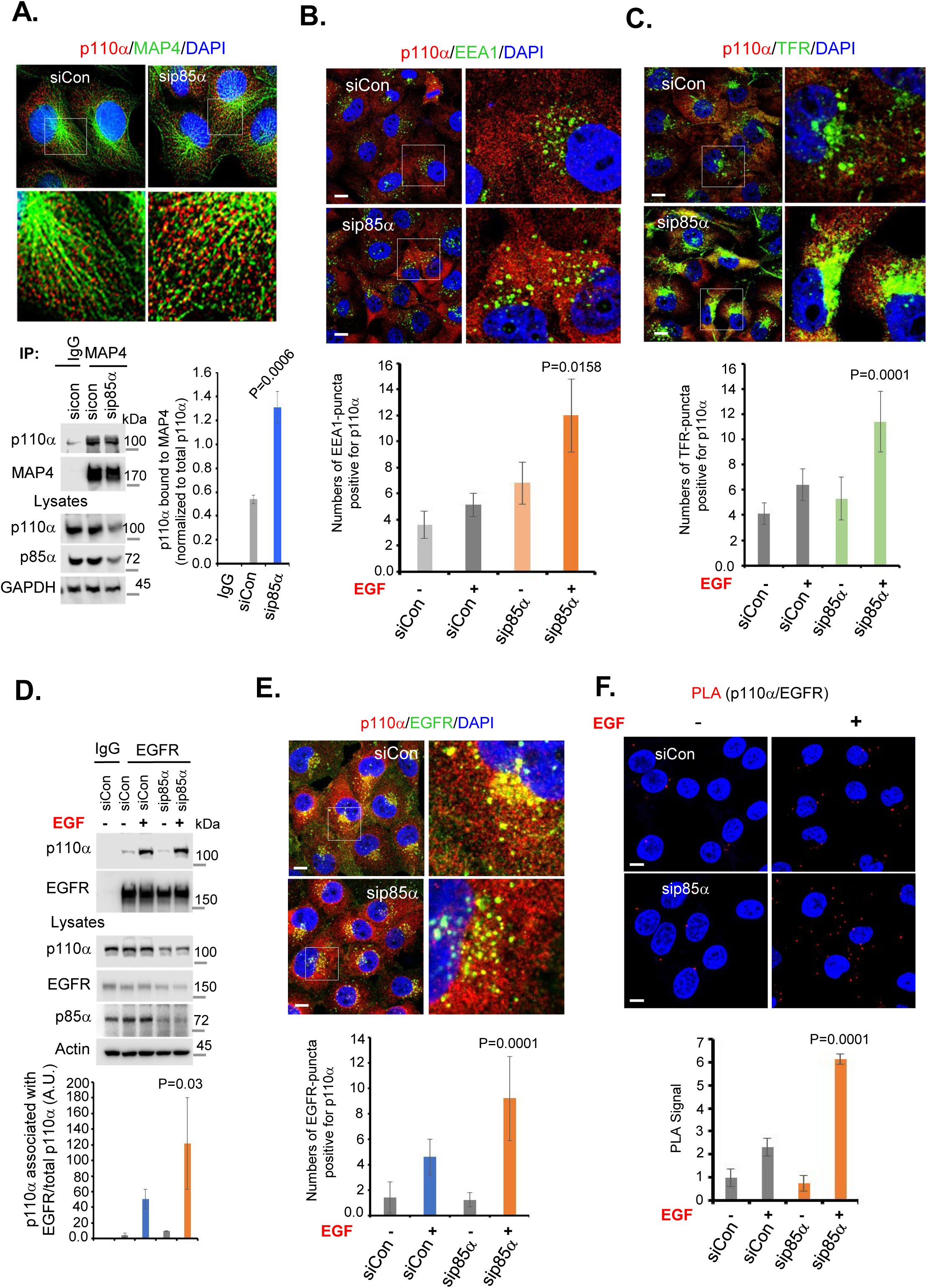
p110α Distribution along Microtubules, Integration into Endosomal Membranes and Association with Agonist-stimulated Receptor Tyrosine Kinases is Enhanced upon p85α KD. **A, p110α localizes along microtubules and associates with MAP4 upon p85α KD.** Upper panel, MDA-MB-231 cells were transfected with control siRNAs or siRNAs targeting p85α. 48-72 hours post-transfection, cells were fixed with 4% PFA. The distribution of p110α along microtubules in cells treated with control and p85α siRNAs was examined by IF using antibodies specific for p110α (red) and MAP4 (green). Similarly, the agonist-stimulated association of residual p110α with MAP4 upon p85α KD by co-IP. The data represent the mean ± SD from three independent experiments. The statistical significance is indicated by p-value (siCon versus p85α siRNA). **B, C, p110α co-localizes with endosomal markers upon p85α KD.** MDA-MB-231 cells were transfected with control siRNAs or siRNA targeting p85α. 48-72 hours post-transfection, cells were stimulated with EGF and fixed with 4% PFA. The co-localization of p110α with endosomal markers in cells treated with control and p85α siRNAs was examined by IF using antibodies specific for p110α (red) and endosomes (EEA1 and TFR) (green). The number of EEA1- and TFR-puncta positive for p110α was counted in at least 20-30 cells. The statistical significance is indicated by p-value (control versus p85α KD, EGF stimulated). **D, EGF-stimulated p110α association with EGFR upon p85α KD.** MDA-MB-231 cells were transfected with control siRNAs or siRNAs targeting p85α. 48-72 hours post-transfection, cells were stimulated with EGF before harvesting the cells. EGFR was IPed using anti-EGFR antibody beads, and co-IPed p110α examined by WB. The data represent the mean ± SD from three independent experiments. The statistical significance is indicated by p-value (siCon versus p85α siRNA, EGF stimulated). **E, EGF-stimulated p110α co-localization with EGFR upon p85α KD.** MDA-MB-231 cells were transfected with control siRNAs or siRNAs targeting p85α. 48-72 hours post-transfection, cells were stimulated with EGF for 5 minutes before fixation with 4% PFA. The co-localization of p110α with EGFR was examined by IF study using antibodies specific for p110α (red) and EGFR (green). EGFR-puncta positive for p110α were counted in at least 20-30 cells. The statistical significance is indicated by p-value (control versus p85α KD, EGF stimulated). **F, EGF-stimulated p110α association with EGFR upon p85α KD by PLA.** MDA-MB-231 cells were transfected with control siRNAs or siRNAs targeting p85α. 48-72 hours post-transfection, cells were stimulated with EGF for 5 minutes before fixation with 4% PFA. Cells were incubated with p110α and EGFR-specific primary antibodies followed by PLA assay. PLA puncta were counted in at least 20-30 cells. The statistical significance is indicated by the p-value (control versus p85α KD, EGF stimulated).

Next, we examined the interaction of p110α with activated receptor tyrosine kinases following p85α KD. Activated receptor tyrosine kinases undergo rapid internalization, and the endosomal compartments are a major platforms for intracellular signaling^26^. As agonist-stimulation was required to drive activation of PI3K/Akt signaling in response to p85α KD, we envisioned that the residual p110α binds activated receptor tyrosine kinases to regulate the agonist-activated PI3K/Akt signaling. Notably, the association of p110α with EGFR was enhanced by EGF stimulation following p85α KD as demonstrated by IF, co-IP, and proximity ligation assays (**Fig. 4D-F**). The increased association of p110α with MAP4 and its distribution along microtubules facilitates the enhanced association of residual p110α, presumably coupled with p85β, with endosomal EGFR in response to EGF stimulation and p85α KD. These results concord with our recent report regarding the spatial localization of PI3K/Akt signaling predominantly in endosomal compartments downstream of activated receptor tyrosine kinases^19^.

### The p110α C2 Domain-PI3P Interaction Promotes Agonist-Stimulated Endosomal PI3K/Akt Signaling in Response to p85α KD

We next examined the mechanisms by which p110α is recruited to the endosomal compartments following agonist-stimulation and p85α KD. The C2 domain of p110α is a presumed membrane-interacting module in PI 3-kinases^24^ that binds the iSH2 domain of p85α by contact at two different sites^27–29^. Moreover, the p85α iSH2 domain partially masks the p110α C2 domain by creating steric hindrances with an anionic phospholipid membrane surface^12, 30^, thereby inhibiting its interaction with membranes^27^. This suggests p85α loss may enhance the p110α C2 domain interaction with membranes to regulate PI3K/Akt signaling. Consistent with this idea, ectopically expressed HA-tagged p110α with the C2 domain deleted disrupted its co-localization with the endosomal marker EEA1 compared to wild-type p110α (**Fig. 5A, Suppl. Fig. 5A**). Strikingly, the C2 domain deletion mutant p110α did not enhance agonist-stimulated Akt activation following p85α KD (**Fig. 5B)**. Furthermore, HA-tagged wild-type p110α, but not the C2 domain deletion mutant p110α (C2 Del HA-p110α), rescued the effects of p110α KD on agonist-stimulated Akt activation (**Fig. 5C**). These results indicate the C2 domain of p110α is required for its recruitment to endosomal membranes and for enhancing agonist-stimulated Akt activation in response to p85α KD.

**Figure 5:**
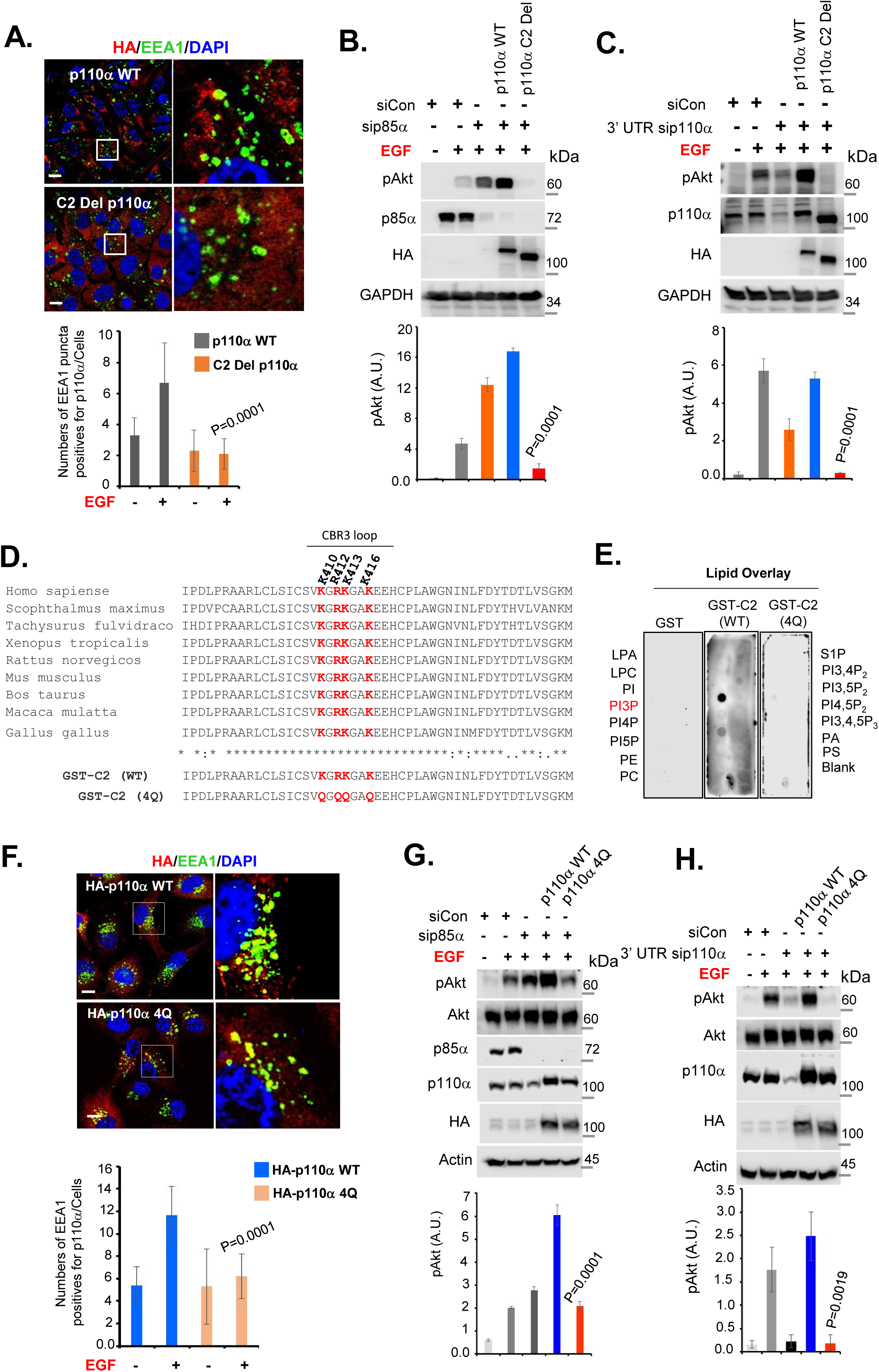
p110α Integration into Endosomes and Agonist-Stimulated PI3K/Akt Signaling upon p85α KD is Regulated by the C2 Domain-PI3P Interaction. **A, deletion of p110α C2 domain impairs its integration into endosomal membranes upon agonist stimulation.** MDA-MB-231 cells stably expressing wild-type HA-p110α or C2 domain deletion HA-p110α were transfected with control siRNAs or siRNAs targeting endogenous p85α. 48-72 hours post-transfection, cells were stimulated with EGF for 5 minutes before fixation with 4% PFA. The cells were immunostained with antibodies for HA (red) and EEA1 (green). EEA1-punta positive for HA were counted in least 20-30 cells. The statistical significance is indicated by p-value (control versus p85α KD, EGF stimulated). **B, C, C2 domain deletion mutant p110α disrupts EGF-stimulated Akt activation upon p85α KD.** B, MDA-MB-231 cells stably expressing empty vector, wild-type HA-p110α or C2 domain deletion HA-p110α were transfected with control siRNAs or siRNA stargeting endogenous p85α. 48-72 hours post-transfection, cells were stimulated with EGF for 5 minutes before harvesting cells. phospho-Akt, p85α and HA-p110α (wild-type or C2 domain deletion mutant) were examined by WB. The data represents the mean ± SD from three-independent experiments. **C,** MDA-MB-231 cells stably expressing empty vector or HA-p110α (wild-type or C2 domain deletion) were transfected with control siRNAs or siRNAs targeting endogenous p110α at 3’ prime UTR regions. 48-72 hours post-transfection, cells were stimulated with EGF for 5 minutes before harvesting cells. Phospho-Akt, endogenous p110α and HA-p110α (wild-type or C2 domain deletion mutant) levels were examined by WB. The data represent the mean ± SD from three independent experiments. The statistical significance is indicated by p-value (wild-type p110α versus C2 domain deletion mutant p110α, EGF stimulated). **D, alignment of the p110α C2 domain from different species.** The conserved lysine and arginine residues in the CBR3 loop are indicated in red. Lower panel, all three lysine and one arginine residues were mutated to glutamine (4Q) to generate the mutant GST-fusion protein of the C2 domain. **E, lipid overlay assay.** The lipid overlay assay was performed by incubating the PIP-Strip membrane with purified GST-fusion proteins (each 0.5 µg/ml) overnight at 4 C°, followed by the detection of bound protein using HRP-conjugated anti-GST antibody. Representative data of three independent experiments were shown. **F, 4Q mutant p110α exhibits impaired integration into early endosomes.** MDA-MB-231 cells stably expressing HA-p110α (WT) or C2 domain mutant p110α (4Q) were stimulated with EGF before fixing the cells with 4% PFA. The cells were immunostained with antibodies for HA (red) and EEA1 (green). EEA1 puncta that co-stained with HA were counted in least 20-30 cells. The statistical significance is indicated by p-value (p110α WT versus p110α 4Q, EGF stimulated). **G, H, the p110α 4Q mutant exhibits impaired EGF-stimulated Akt activation upon p85α kD.** MDA-MB-231 cells stably expressing empty vector or HA-p110α (WT of 4Q mutant) were transfected with control siRNAs, p85α siRNAs (G) or siRNAs targeting endogenous p110α at 3’ prime UTR region (H). 48-72 hours post-transfection, cells were stimulated with EGF for 5 minutes before harvesting cells. phospho-Akt, endogenous p110α and p85α, and HA-p110α WT or 4Q mutant levels were examined by WB. The data represent mean ± SD from three independent experiments. The statistical significance is indicated by p-value (p110α WT versus p110α 4Q, EGF stimulated).

To define the critical amino acid residues in the C2 domain of p110α, we mutated the basic residues K410, R412, K413 and K416 in the CBR3 loop of C2 domain that are the putative membrane-interacting residues^27^ and are highly conserved in mammals (**Fig. 5D**). The C2 domain assumes a characteristic eight-stranded antiparallel β-sandwich^27^. We performed a lipid overlay assay using the GST-tagged C2 domain and its 4Q mutant form (K410, R412, K413 and K416 each mutated to Q). The p110α C2 domain, but not its 4Q mutant, bound robustly to PI3P, a critical endosomal phosphoinositide^14^. Additionally, HA-tagged 4Q mutant p110α exhibited impaired endosomal localization (**Fig. 5F; Suppl. Fig. 5B**). These mutations severely abrogated an ability of p110α to support agonist-stimulated Akt activation following p85α KD (**Fig. 5G, H**). Furthermore, these mutations also greatly disrupted the association of p110α with EGFR and the co-localization with endosomal EGFR upon EGF stimulation (**Fig. 6A, B, C; Suppl. Fig. 6A**). The impaired association of 4Q mutant p110α with EGFR receptor upon agonist stimulation was further validated by co-immunoprecipitation and PLA after transient transfection (**Fig. 6D, E, F**). Together, these data indicate that the p110α C2-PI3P interaction is pivotal for bringing p110α to the endomembrane to associate with activated receptor. The loss or reduced expression of p85α leads to enhanced integration of p110α into endosomal membranes and its interaction with activated receptor, leading to increased agonist-stimulated PI3K/Akt signaling (**Fig. 6G**).

**Figure 6:**
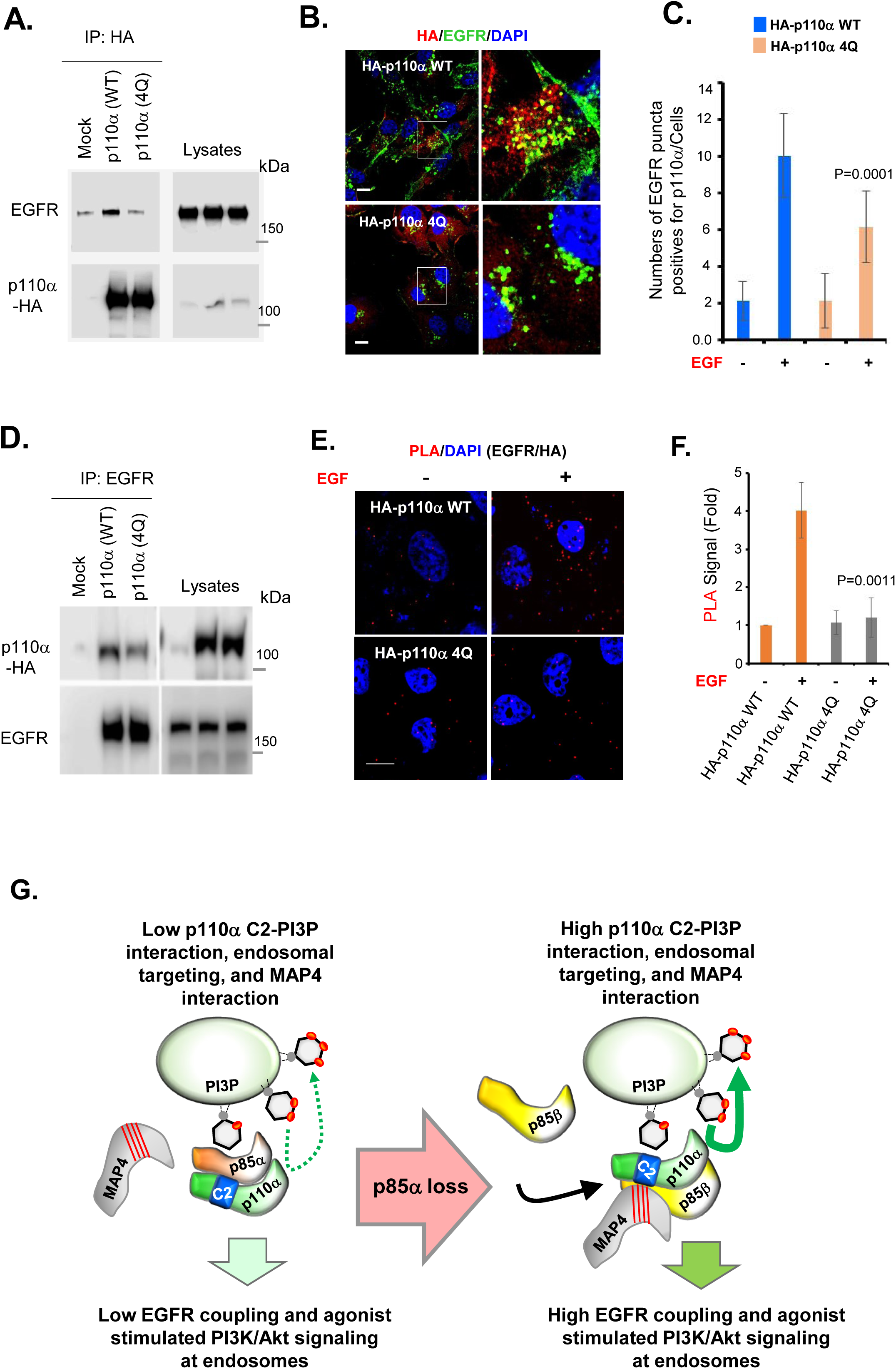
p110α Association with Agonist-stimulated Receptor Tyrosine Kinase (For this experiments, p85α KD was not used)is Regulated by the C2 Domain-PI3P Interaction. **A,** MDA-MB-231 cells stably expressing HA-tagged WT p110α or 4Q mutant p110α were stimulated with EGF for 5 minutes before harvesting the cells. p110α was IPed using anti-HA antibody agarose beads and the co-IPed endogenous EGFR was examined by WB. The data is representative of three independent experiments. **B, C,** MDA-MB-231 cells stably expressing HA-tagged WT or 4Q mutant p110α were stimulated with EGF for 3-5 minutes before fixing the cells with 4% PFA. The cells were immuno-stained with antibodies for HA (red) and EGFR (green). The EGFR-puncta that co-localized with HA were counted in least 20-30 cells. The statistical significance is indicated by p-value (p110α WT versus p110α 4Q, EGF stimulated). **D,** COS-7 cells transiently expressing HA-tagged WT p110α or 4Q mutant p110α were stimulated with EGF for 5 minutes before harvesting the cells. Endogenous EGFR was IPed and the co-IPed p110α was examined by WB. The data is representative of three independent experiments. **E, F,** COS-7 cells transiently expressing HA-tagged WT or 4Q mutant p110α were stimulated with EGF for 3-5 minutes before fixing the cells with 4% PFA. The cells were processed for PLA assay. The PLA punta were counted in least 20-30 cells and normalized to control cells. The statistical significance is indicated by p-value (p110α WT versus p110α 4Q, EGF stimulated). **G,** Schematic diagram depicting the mechanism on how loss of p85α adaptor subunit promotes agonist stimulated PI3K/Akt signaling in the endomembrane. The loss of p85α adaptor subunit promotes the spatial localization of residual p110α catalytic subunit along microtubules via MAP4 binding and integration into endosomal membranes via p110α C2 domain and endosomal PI3P interaction. This leads to increased interaction of residual p110α catalytic subunit with activated endosomal receptor, promoting PI3K/Akt signaling in the endosomal membrane.

## DISCUSSION

We recently reported that agonist-stimulated PI3,4,5P_3_ generation and Akt activation occur predominantly at internal membranes^19^. The rapid endocytosis of activated receptor tyrosine kinases and PI3Kα organization along microtubules via MAP4 promotes spatial activation of PI3K/Akt signaling at the endomembrane by facilitating the association of PI3Kα with activated receptor tyrosine kinases^19^. In the current study, we deciphered the comprehensive mechanism by which loss of the p85α adaptor subunit paradoxically elicits enhanced PI3K/Akt signaling downstream of activated receptor tyrosine kinases by emphasizing the inherent capacity of p110α catalytic subunit for membrane interaction. The organization of p110α along microtubules, p110α association with receptor tyrosine kinase and p110α recruitment into internal membrane compartments remain completely intact upon p85α KD and are augmented compared to basal levels when normalized for the reduction in p110α levels. Notably, the p110α C2 domain interaction with PI3P guides the spatial recruitment of p110α into endosomes, where the majority of receptor tyrosine kinase remains active after agonist stimulation. As a result, p110α elicit pronounced PI3K/Akt signaling downstream of activated receptor tyrosine kinases in response to p85α loss.

p85α and p85β constitute virtually all the adaptor subunits (⁓97%) of class IA PI3K with no detectable expression of p55γ in the cell lines used in these experiments. We observed that in response to p85α KD, the majority of p110α associates with p85β. Moreover, p85β KD abrogates the effects of p85α KD in augmenting agonist-stimulated PI3K/Akt signaling, underscoring the key functional role of p85β in the paradoxical PI3K/Akt hyperactivation upon p85α loss. Consistent with these results, p110α coupled with p85β was reported to be more competent at inducing PI3K/Akt signaling^12^. Furthermore, p110β (i.e., PI3Kβ) cannot assume the predominant role in receptor tyrosine kinase stimulated PI3K/Akt signaling even in p85α-p110α downregulated condition because of the more stringent constraint imposed on p110β by p85 adaptor subunits^31^. Although p110α devoid of p85α and expressed in mammalian cells is enzymatically inactive as opposed to in insect or yeast cells as demonstrated by *in vitro* kinase assay^32, 33^, the residual p110α uncoupled with p85 may also contribute to agonist-stimulated PI3K/Akt signaling upon p85α loss in mammalian cells. It seems plausible that upon p85α loss, residual p110α remains active due to disengaged intermolecular constraints imposed by the p85α nSH2 and iSH2 domains. However, unlike p110α helical domain mutants (*e.g.*, E542K; E545K) which disrupt the p85 nSH2 domain interaction and induce the constitutive activation of PI3K/Akt signaling independent of receptor tyrosine kinase activation^34^, agonist stimulation is required for the enhanced activation of PI3K/Akt signaling in response to p85α KD. This finding indicates that rapidly internalized active receptor tyrosine kinases in endosomes are indispensable for eliciting the spatial activation of p110α at endomembranes.

By deleting and mutating key basic residues in the C2 domain of p110α, we have demonstrated that the interaction of this domain with the phosphoinositide PI3P is required for its recruitment to endosomal membranes and activating agonist-stimulated PI3K/Akt signaling upon p85α KD. These findings are consistent with the observation that the p85α iSH2 domain partially masks the C2 domain from interacting with membranes^27^. Additionally, the C2 domain mutant p110α (N345K) or deletion of the C-terminal segment of the C2 domain, which disrupts the p85α iSH2 domain interaction, promotes PI3K/Akt signaling^29^. Furthermore, the more negative electrostatic charge distribution in the p85α verus the p85β iSH2 domain creates a steric hindrance for membrane interaction^12^. Collectively, these studies support a model whereby the p110α C2 domain is not fully exposed for PI3P binding and endosomal recruitment until hindrance from p85α is removed^19^. Upon p85α loss, the resulting enhanced C2 domain-mediated recruitment of p110α to endomembranes promotes its interaction with receptor tyrosine kinases activated by agonist stimulation, resulting in augmented PI3K/Akt signaling. As many components of PI3K/Akt signaling including activated receptor tyrosine kinases, PI3K, Akt and mTOR are localized in endosomes and internal membrane compartments^35^, therapeutic targeting of the molecular interaction of the p110α C2 domain with PI3P could prove to be a promising strategy to disrupt receptor tyrosine kinase stimulated endosomal PI3K/Akt signaling in cancer, including those with loss or diminished expression of p85α. Indeed, a recent report of an allosteric PI3Kα activating compound ^36^ indicates that its binding diminishes p85α interactions with the p110α kinase domain and the inhibitory interface between p85α and the p110α-C2 domain, which would be predicted to enhance p110α interaction with the lipid membrane and PI3P. This mechanism of activation of PI3Kα is fully consistent with our model for p85α/β, MAP4 and PI3P regulation of PI3Kα (Fig. 6G).

## ACKNOWLEDGMENTS

We thank members of the Anderson and Cryns lab for comments and Dr. Adrea Galmozzi for discussions and comments, and Lance Rodenkirch for technical support. This work was supported by a National Institutes of Health grant R35GM134955 (R.A.A.), National Institutes of Health grant 5R21AG074605-02 (N. T. and R. A.A.), Department of Defense Breast Cancer Research Program grants W81XWH-17-1-0258 (R.A.A.), W81XWH-17-1-0259 (V.L.C.) and W81XWH-21-1-0129 (V.L.C.), and a grant from the Breast Cancer Research Foundation (V.L.C.).

## AUTHORS CONTRIBUTIONS

N.T., M.C., V. C., and R.A.A. designed and discussed experiments. N.T. and M.C. performed experiments. N.T., M.C., V. C., and R.A.A. discussed and wrote the manuscript.

## CONFLICT OF INTEREST

The authors declare no competing interests.

## Methods

### Antibodies

Rabbit anti-p110α (#4249, Cell Signaling), rabbit anti-p110α (#ab40776, Abcam), rabbit anti-p110β (#3011, Cell Signaling), rabbit anti-p85α (#ab191606, Abcam), mouse anti-p85β (#MAB6777, R and D), mouse anti-PI3,4,5P_3_ antibody (Z-P345, Echelon Biosciences), rabbit anti-Akt (#ab126811, Abcam), rabbit anti-Akt-HRP (#4298, Cell Signaling), rabbit anti-phospho-Akt T308 (#2965, Cell Signaling), rabbit anti-phospho-Akt S473 (#4060, Cell Signaling), rabbit anti-pAkt (OMA1-03061, Life Technologies), rabbit anti-phospho Akt (#OMA1-03061, Invitrogen), mouse anti-phospho Akt (#200-301-268S, Rockland), mouse anti-MAP4 (#sc-390286, Santa Cruz), rabbit anti-MAP4 (#A301-488A, Bethyl), mouse anti-MAP4 agarose beads (#sc-390286 AC, Santa Cruz), normal mouse IgG beads (sc-2343, Santa Cruz), rabbit anti-HA (#3724, Cell Signaling), mouse anti-HA agarose beads (#26181, ThermoFisher), anti-GST HRP conjugate (#RPN 1236V, Amersham), rabbit anti-Flag (#2368, Cell Signaling), mouse anti-Flag (#F1804, Sigma), anti-Flag M2 affinity gel (#F2426, Sigma-Aldrich), rabbit anti-GAPDH (#2118, Cell Signaling), rabbit anti-tubulin (#ab18251, Abcam), mouse anti-EEA1 (#610457, BD Biosciences), rabbit anti-EEA1 (#3288, Cell Signaling), mouse anti-TFR (#136800, Invitrogen), rabbit anti-TFR (#ab84036, Abcam), rat anti-tubulin (#ab6160, Abcam), rabbit anti-tubulin (#ab6046, Abcam), normal rabbit IgG beads (#2729, Cell Signaling), rabbit anti-EGFR beads (#5735 Cell Signaling), rabbit anti-EGFR (sc-03, Santa Cruz), mouse anti-EGFR (#sc-120, Santa Cruz), anti-beta actin (#4967), rabbit anti-Na+/K+ ATPase (#3010, Cell Signaling) and PTEN (sc-7974, Santa Cruz).

### Cell Culture

MDA-MB-231, A431, H4, HEK293T, HCT116 and COS-7 were purchased from American Type Culture Collection (ATCC). All cell lines were cultured in DMEM-containing 10% FBS (Hyclone) and antibiotics (penicillin/streptomycin) at 37°C in a 5% CO_2_ incubator. All cell lines were routinely tested for mycoplasma contamination and sub-cultured before 80-90% confluency. For examining the Akt activation or PI3,4,5P_3_ generation by FBS (10%) or EGF (10 ng/ml) or insulin (10 ng/ml), cells were serum-starved overnight before stimulation as described previously^19, 37^.

### Immunoprecipitation and Immunoblotting

For immunoprecipitation, the cells were often grown and harvested from 10 cm culture dishes. Before cell lysis, cells were washed with cold 1X PBS 2 times. Cells were lysed using lysis buffer (50 mM Tris-HCl pH 7.4, 150 mM, 1% Triton-X100, 1 mM EDTA, 1 mM EGTA and protease/phosphatase inhibitors). The clear supernatants were obtained by centrifuging at 14,000 rpm at 4°C. Clear supernatants were incubated with antibody-coated agarose beads or control beads overnight at 4°C. If nascent antibodies were used, protein-antibody complexes were isolated using protein G or A Sepharose 4B beads (Amersham). Beads were washed three times with lysis buffer before eluting the immunocomplexes with 2x sample buffer, run through SDS-PAGE gel and subjected to immunoblotting using specific antibodies. For immunoblotting/western blotting, primary antibodies were diluted 1:2000 in 3% BSA in TBS-T (0.1% Tween 20). This was followed by incubation with HRP-labelled secondary antibodies. The images were acquired using LI-COR Odyssey FC.

### Small Interfering RNA (siRNA)

Control siRNA (siCon):5’-UUUCCGCACUGUGAUUCGG-3’

sip85α I: 5’-GGAUCAAGUUGUCAAAGAA-3’

sip85α II: 5’-GCAGCUGAGUAUCGAGAAA-3’

sip85α III: 5’-GGGTGACATATTGACTGTGAATAAA-3’

siPTEN: 5’-GCUUGAAGACUAAAGCAUA-3’

sip110α: 5’-UCAAGAAGAAAGCUGACCAUGCUGC-3’

sip110β: 5’-GCUUCAGAUUUGGCCUAAA-3’

siMAP4: 5’-CCGGGAACUCAGAGUCAAA-3’

These siRNA oligonucleotides were designed using Invitrogen Block-iT RNAi Designer and purchased from Thermo Fisher or Dharmacon.

### siRNA or Plasmid Transfection or Lentiviral Infection

For transfection of siRNA, LipofectamineRNAiMAX (#13778150, Invitrogen) was used following the protocol provided by the manufacturer, and cells were assayed 48-72 hours post-transfection. Lipofectamine-3000 (#L3000015, Invitrogen) was used for transient transfection of plasmids, and cells were harvested 24-48 hours post-transfection. For the stable expression of HA-tagged wild type p110α or its mutant forms (C2 deletion mutant, HA-p110α C2 Del., or p110α (4Q)), the pWPT-GFP lentiviral vector system was used as described previously ^19, 38, 39^.

### Immunofluorescence (IF) Staining and Confocal Microscopy

For immunofluorescence study, cells were grown on glass coverslips. Cells were fixed with 4% PFA followed by permeabilization with 0.1% Triton-X100 and blocking with 3% BSA in TBS. Cells were incubated with a primary antibody overnight at 4°C followed by incubation with fluorescent-conjugated secondary antibodies (Molecular Probes) for 1 hour at room temperature. Cells were mounted in Prolong^TM^ Glass Antifade Mounting media (#P36984, Thermo Fisher Scientific). The images were taken by Leica SP8 3X STED Super-Resolution Microscope, which is both a point scanning confocal and 3X STED super-resolution microscope. The Leica SP8 3X STED microscope was controlled by LAS-X software (Leica Microsystems). Images were acquired using 60x or 100x objective lens. Only the image in Figure 4A were taken by Nikon TE2000-U and image processed using Metamorph. For quantification of fluorescence intensity, the mean fluorescent intensity of interested channels in each cell (at least 20 cells used) was measured by LAS-X. The images were processed using Image J.

For the investigation of activated Akt or PI3,4,5P_3_ lipid messenger generated by immunofluorescence study, the cells growing in the coverslips were stimulated with EGF stimulation followed by rapid fixation with 4% PFA prepared in TBS containing the phosphatase inhibitors (2.5 mM NaF and 2.5 mM Na3VO4). For the immunofluorescence staining of MAP4, cells were fixed in ice-cold methanol and anti-MAP4 antibody (#sc-390286, Santa Cruz) was used. Following 3-times washing with TBS, cells were permeabilized with 0.1% Triton-X100 in TBS-containing phosphatase inhibitors. Then, cells were incubated in the blocking buffer containing 3% BSA in TBS for 1 hour at room temperature followed by overnight incubation with primary antibody (prepared in TBS-T containing 3% BSA) at 4°C in humidified chamber. Cells were washed 3 times with TBS-T (TBS-containing 0.1% Tween 20) followed by incubation with secondary antibody for 1 hour. Cells were washed 3 times before mounting.

### Examination of Activated Akt in Endosomes

For endosome isolation, cells grown in 10 cm culture plates were used. Cells after EGF stimulation, cells were fractionated using an endosome isolation kit (ED-028, Invent Biotechnologies). Briefly, cells were washed with cold TBS on ice before detaching the cells with Trypsin-EDTA. Then, collected cells were suspended in buffer A. The cell supernatant was centrifuged at first to remove the plasma membrane/larger organelles before using supernatant for endosome isolation. The equal amount of isolated plasma membrane/larger organelles vs endosomal fractions were run through SDS-PAGE and immunoblotted with activated Akt and markers for plasma membrane and endosomes.

### Semiquantitative Mass Spectrometry Analysis of p110α Immunocomplex from p85α KD Cells

MDA-MB-231 cells stably expressing the HA-p110α and grown in 15 cm culture plates were used. Cells were transfected with p85α siRNA and used 48-72 hours-post transfection. Cells were lysed using lysis buffer (50 mM Tris-HCl [pH 7.4], 150 mM NaCl, 0.5% Triton-X100, 1 mM EDTA, 10 mM NaF and 5 mM Na3VO4) containing protease/phosphatase inhibitors (Roche). Cells were incubated in lysis buffer for 2-3 hours in rotator at 4°C. After centrifuging at 14,000 rpm for 15 minutes, the clear cell lysates were used to immune-precipitate HA-p110α using HA agarose beads. Isolated immunocomplex were eluted using 2x sample buffer without dye. The immunocomplexes obtained from mock or control siRNA or p85α KD cells were analyzed at the mass spectrometry facility of UW-Madison Biotech Center. The spectral count of the p110α, p85α, and p85β peptides obtained were used for the analysis. However, p55γ was not detected in MDA-MB-231 cells by mass spectrometry analysis.

### Lipid Overlay Assay

PIP Strips membranes (#P-6001, Echelon Biosciences) were blocked with 3% fatty acid free BSA (#A7030, Sigma) in TBS for 1-2 hours at room temperature. After blocking, the membrane was incubated with purified GST-fusion proteins (GST alone, GST-C2 and GST-C2 4Q) diluted in 3 ml TBS-T 3% BSA at the concentration of 0.2 µg/ml. The membrane was washed with 5 ml TBS-T three times with gentle agitation for 10 minutes each. This was followed by incubation with anti-GST-HRP antibody diluted in TBS-T 3% BSA for 1 hour in room temperature. The membrane was washed 3-5 times with TBS-T before examining the bound GST-fusion protein using ECL detection system.

### Proximity Ligation Assay (PLA)

PLA was applied to detect *in situ* protein-protein interaction as described previously^19, 37^. Cells after fixation and permeabilization were blocked before incubation with primary antibodies as in the routine IF staining procedure. After that, the cells were processed for PLA (#DUO92101, Millipore Sigma) according to the manufacturer’s instruction and previously. PLA signals are detected by Leica SP8 confocal microscope as discrete punctate foci and provide the intracellular localization of the protein-protein complex and were later quantified by ImageJ.

### Subcloning of Human p110α and its Constructs into PWPT-GFP Lentiviral Vector

The cDNA for human p110α was a kind gift of Dr. Peter K. Vogt (Scripps Research Institute). The detailed procedure for subcloning of p110α in frame with HA-tag at N-terminus of pWPT-GFP lentiviral vector has been described previously^19^. Similarly, the generation of C2 domain deletion mutant of p110α has also been described previously^19^. For the generation of p110α 4Q mutant, p110α cDNA was cloned, at first into SalI and XhoI sites of pCMV-HA vector using primers: 5’- AAAGTCGACCATGCCTCCACGACCATCATC-3’ and 5’- AAACTCGAGTCAGTTCAATGCATGCTGT-3’. Then, C2 domain residues Lys^410^, Arg^412^, Lys^413^ and Lys^416^ were mutated to Glutamine (Q) in two steps. At first, the mutation on Lys^410^ and Lys^416^ were created using primers: 5’-TGCTCTGTTCAAGGCCGAAAGGGTGCTCAAGAGGAAC-3’ and 5’- GTTCCTCTTGAGCACCCTTTCGGCCTTGAACAGAGCA-3’. This mutant was used as a template to generate remaining mutation in Arg^412^ and Lys^413^ using a second set of primers: 5’- TGCTCTGTTCAAGGCCAACAGGGTGCTCAAGAGGAAC-3’ and 5’- TGCTCTGTTCAAGGCCAACAGGGTGCTCAAGAGGAAC-3’. The generated 4Q mutant p110α in pCMV-HA vector was further subcloned into pWPT-GFP lentiviral vector as described previously^19, 39^. The mutations created and integrity of DNA were confirmed by DNA sequencing.

### Cloning of C2 Domain and its Mutant Form into pGEX-6p-1 Vector and Protein Purification

The PCR products for p110α C2 domain and its 4Q mutant form were generated by using the primers: 5’- AAAGGATCCAGTGCACTCAGAATAAA-3’ and 5’- AAACTCGAGTCATTATAGTCTGTTACTCAGTC-3’. The amplified PCR products were subcloned into BamH1 and XhoI sites of pGEX-6p-1 vector (Amersham). The integrity of DNA sequences was validated by DNA sequencing.

The *E. coli* BL21 (DE3) (Thermo Fisher Scientific) were used for the expression of the GST-fusion proteins. Almost all expressed proteins were recovered in the insoluble fractions (inclusion bodies). To recover proteins in soluble fraction, we induced the expression of each protein at 25°C by culturing the bacteria (2 liters of LB medium for each protein) for 48 hours in the presence of 0.1 mM of isopropyl-β- D-thiogalactoside (IPTG) and osmotic stress (330 mM sorbitol and 2.5 mM betaine) as described previously^19^. Proteins were purified from soluble fractions of the bacterial cell lysates using Glutathione Sepharose beads (GE Healthcare). Proteins were dialyzed in 1x TBS overnight at 4°C overnight and protein quantified. The integrity and purity of purified GST-fusion proteins were examined by running 1 µg of proteins through 12% SDS-PAGE gel followed by Coomassie staining.

### Statistics and Reproducibility

All the raw data are available in Statistics Source Data. Data are presented as mean±SD from at least three independent experiments with similar results. All the micrographs (IF images) are the representative images of three representative experiments as indicated in each figure legend. For the quantification of immunofluorescence images, the number of cells used for each representative experiment is indicated and unpaired t-test was conducted to determine the *p*-value between the two groups. The *p*-value less than 0.05 were considered significant between two groups.

### Resource Availability

All other data supporting the findings of this study are available from the corresponding author upon reasonable request. Source data are provided in this paper. For research materials used in this study is available upon reasonable request to corresponding author.

## SUPPLEMENTARY INFORMATION

**Figure S1:**
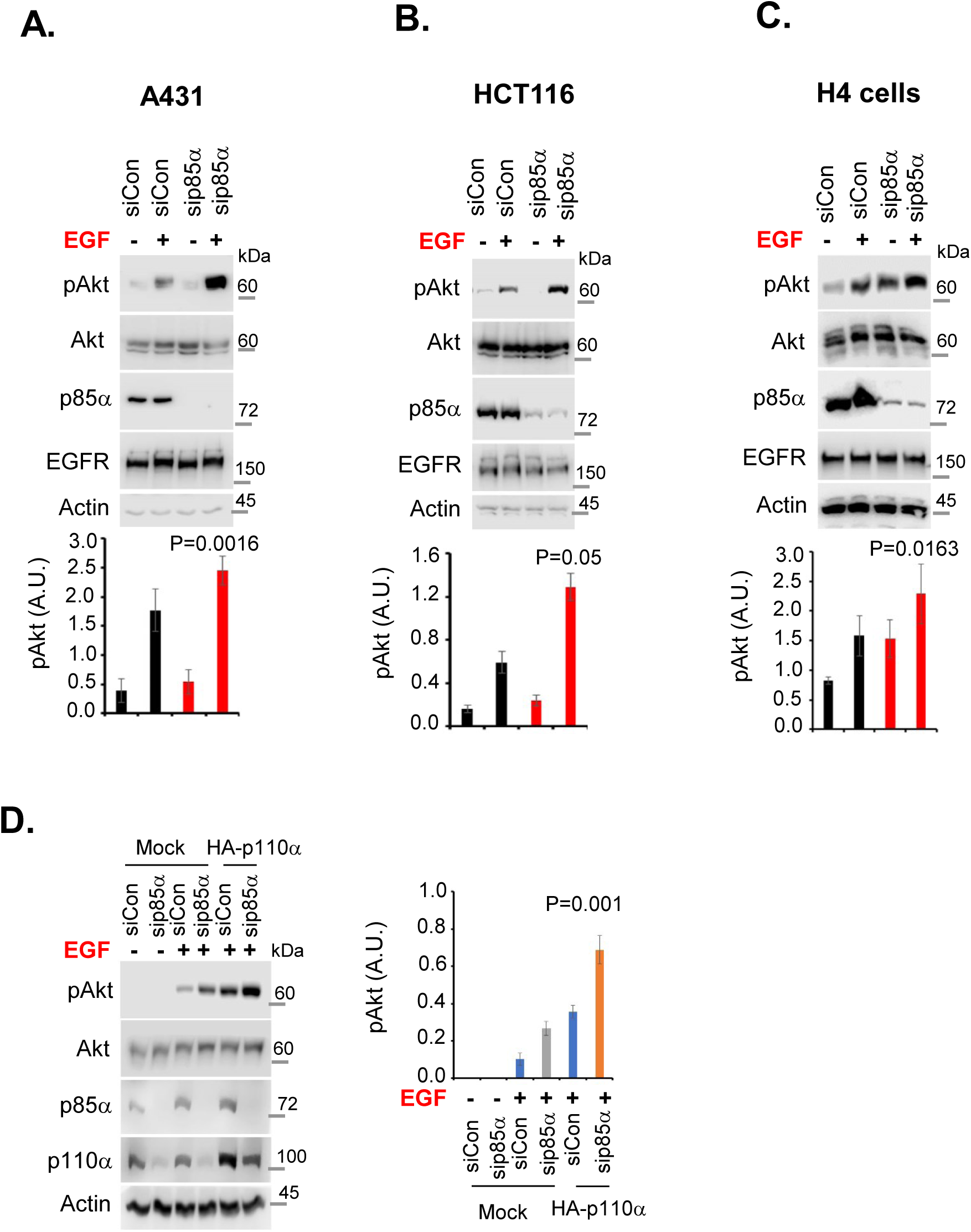
p85α KD Promotes Agonist-stimulated Akt Activation in Multiple Cell Lines. **A, B, C,** A431, HCT116 and H4 cells were transfected with control siRNAs or p85α siRNAs targeting endogenous p85α. 48-72 hours post-transfection, cells were stimulated with EGF for 5 minutes before harvesting the cells. phospho-Akt and p85α levels were analyzed by WB. The data represent the mean ± SD from three independent experiments. The statistical significance is indicated by p-value (siControl versus sip85α, EGF stimulated). D, p85α KD promotes EGF-stimulated Akt activation in p110α overexpressing cells. Mock or HA-p110α overexpressing cells were transfected with siRNAs or p85α siRNAs targeting endogenous p85α. 48-72 hours post-transfection, cells were stimulated with EGF for 5 minutes before harvesting the cells. phospho-Akt and p85α levels were analyzed by WB. The data represent the mean ± SD from three independent experiments. The statistical significance is indicated by p-value (sip85α in mock vs p110α overexpressing cells upon EGF stimulation). Related to Main Figure 1

**Figure S2.**
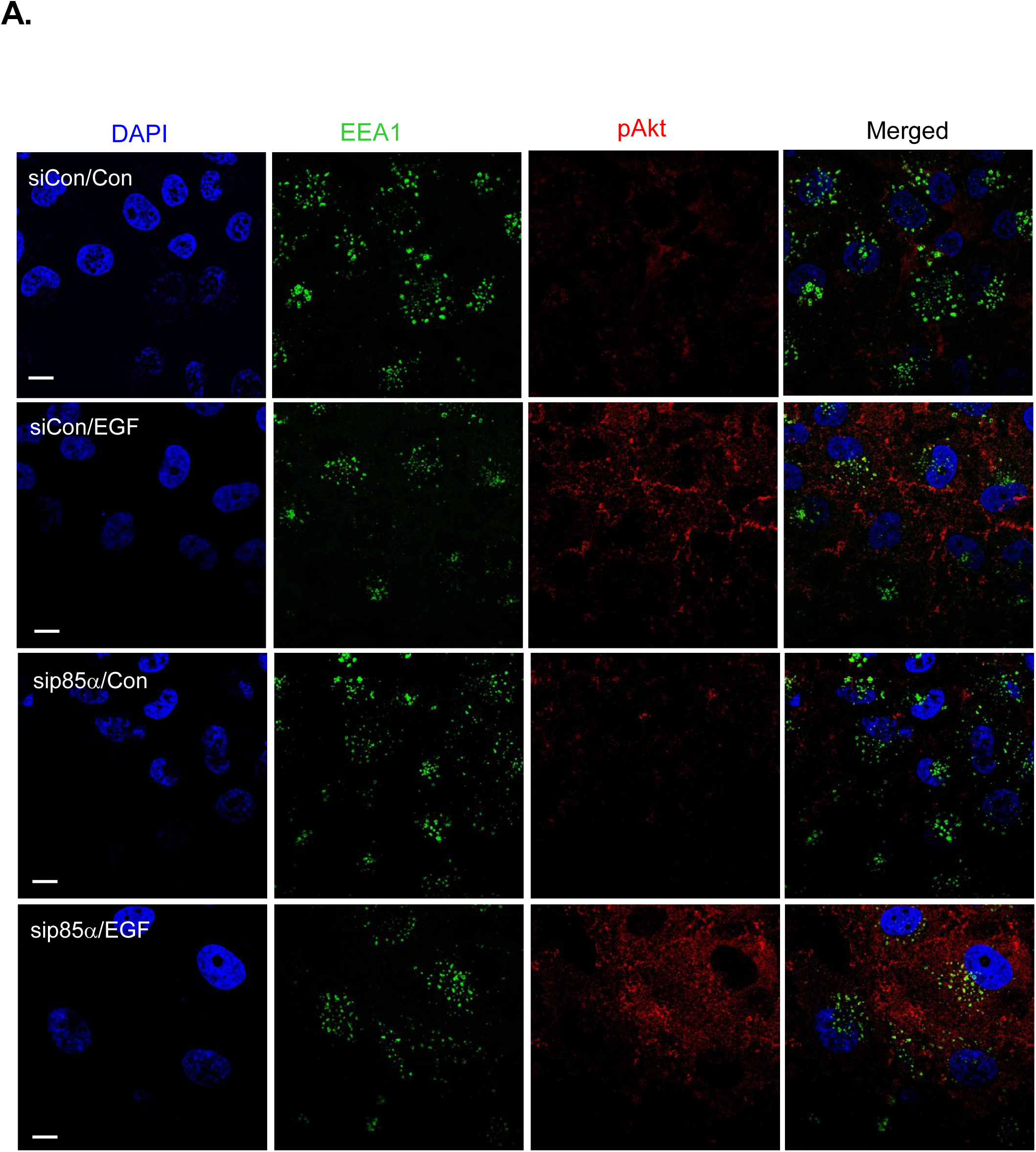
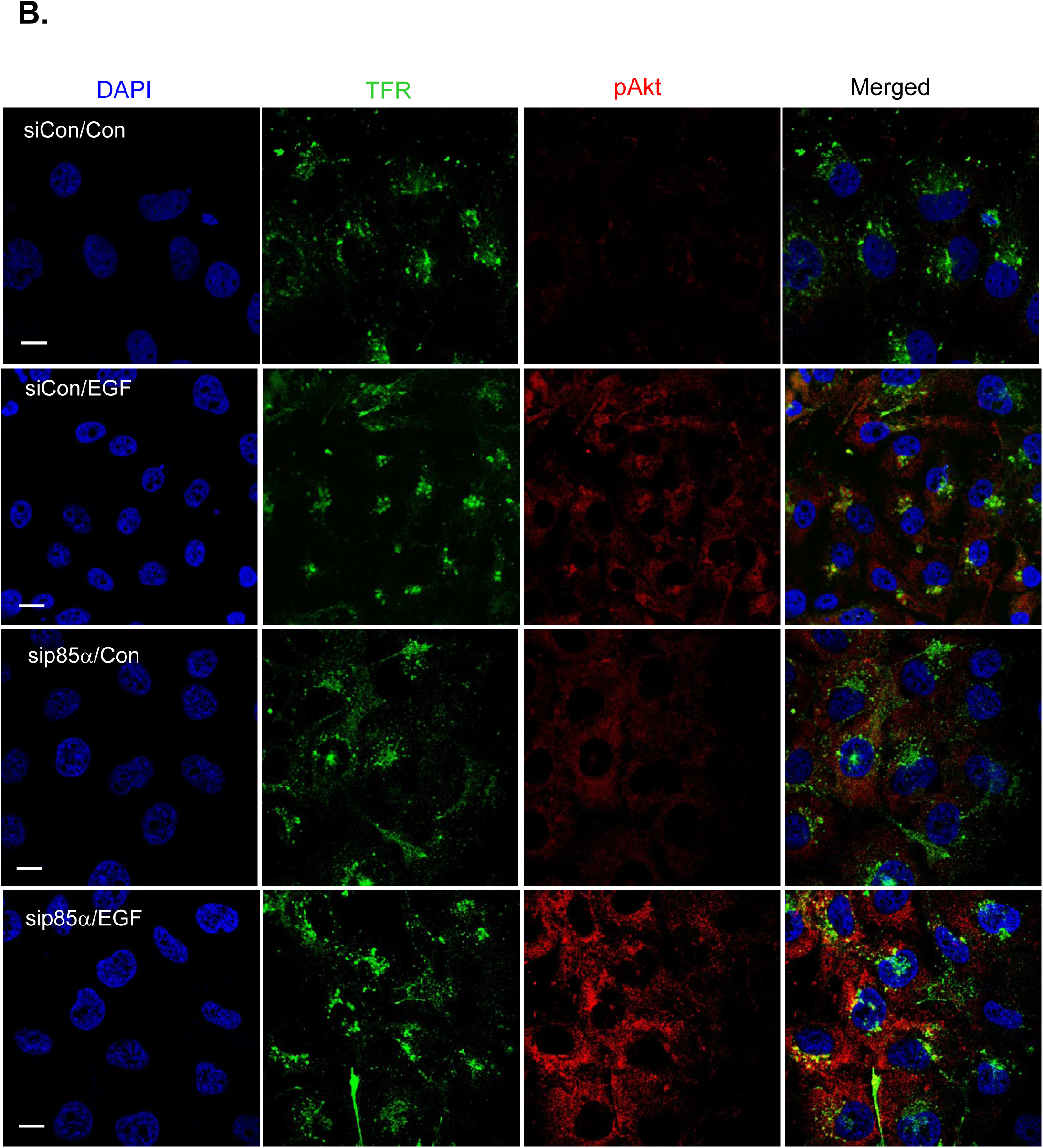
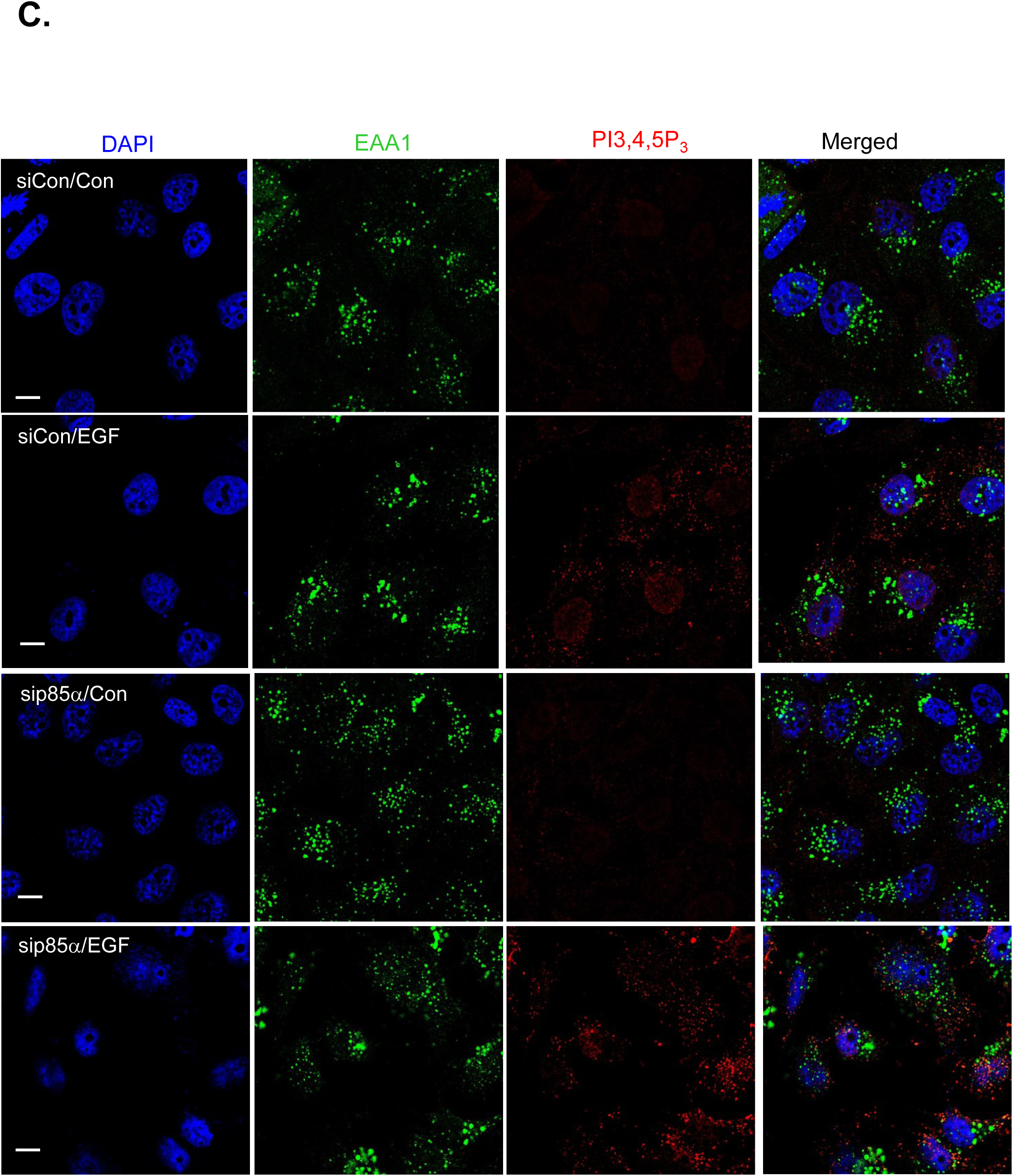
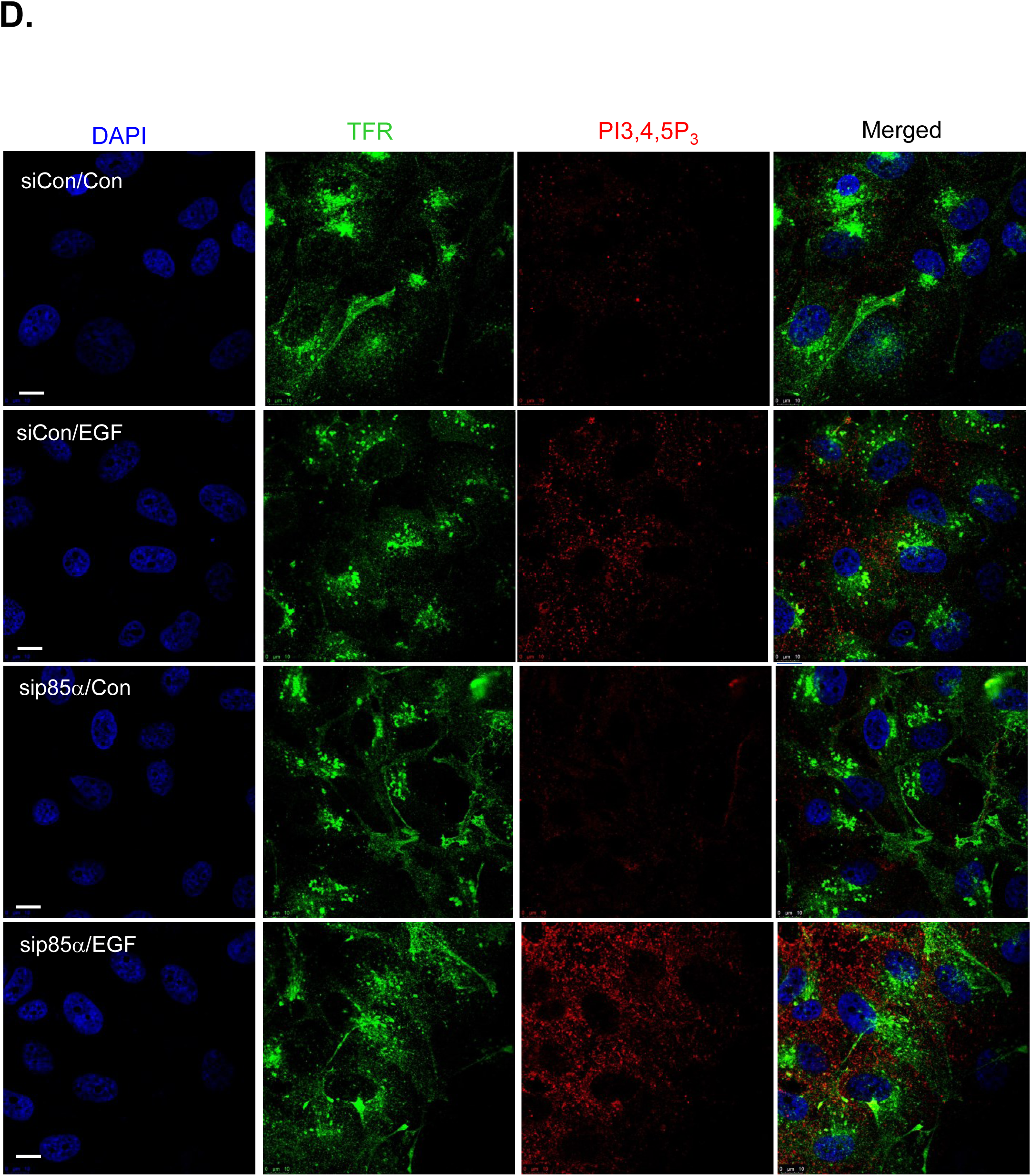
EGF stimulated PI3,4,5P_3_ generation and Akt activation in the endosomes upon p85α KD. **A, B, EGF stimulated Akt activation at endosomes.** MDA-MB-231 cells were transfected with either control siRNA or siRNA targeting p85α. 48-72 hours post-transfection, cells were stimulated with EGF before cells fixation with 4% PFA. The EEA1- and TFR-puncta (green) co-localized with activated Akt (red) were counted in at least 20-30 cells. The images shown are representative images; part of these are shown in Figure 2 (D, E). **C, D, EGF stimulated PI3,4,5P_3_ at endosomes upon p85α KD.** MDA-MB-231 cells were transfected with either control siRNA or siRNA targeting p85α. 48-72 hours post-transfection, cells were stimulated with EGF before cells fixation with 4% PFA. The EEA1- and TFR-puncta (green) co-localized with PI3,4,5P_3_ (red) were counted in at least 20-30 cells. The images shown are representative images; part of these are shown in the main Figure 2 (F, G). Related to Main Figure 2

**Figure S3.**
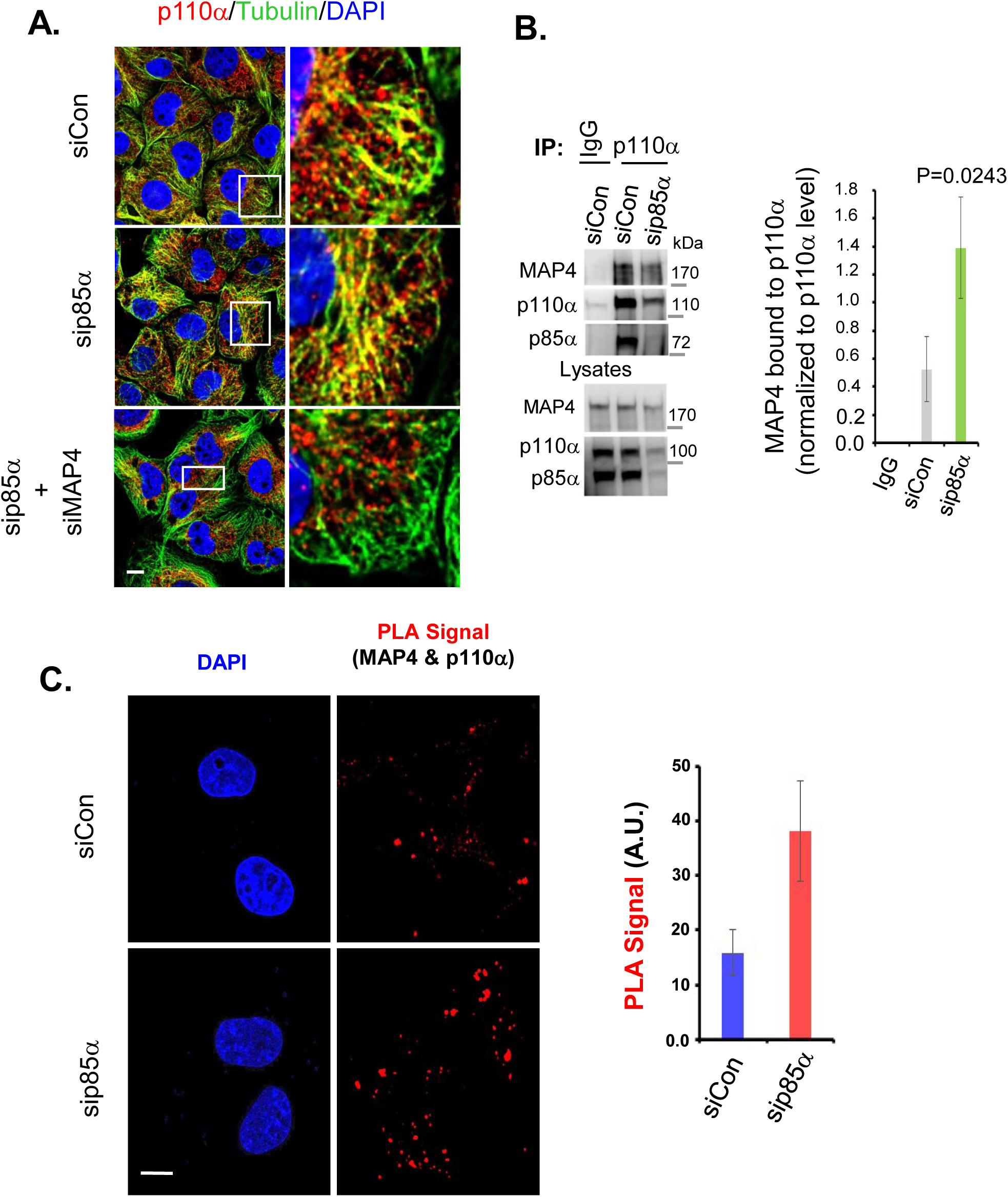
Enhanced p110α Catalytic Subunit Interaction with MAP4 upon p85 KD. **A, MAP4 KD impairs p110α distribution along microtubules.** MDA-MB-231 cells were transfected with control siRNAs, p85α siRNAs or combination of p85α and MAP4 siRNAs. 48-72 hours post-transfection, cells were fixed with methanol. The co-localization of p110α catalytic subunit with microtubules was examined by IF using antibodies specific to p110α (red) and beta-tubulin (green). **B, increased association of p110α with MAP4 upon p85α KD.** MDA-MB-231 cells were transfected with control siRNAs or p85α siRNAs. 48-72 hours post-transfection, cells were harvested and endogenous p110α IPed and the MAP4 co-IPed with p110α was examined by WB. The data represent the mean ± SD from three independent experiments. The statistical significance is indicated by p-value (siControl versus sip85α, EGF stimulated). **C, examination of p110α association with MAP4 by PLA.** MDA-MB-231 cells were transfected with control siRNAs or siRNA stargeting p85α. 48-72 hours post-transfection, cells were fixed with 4% PFA and processed for PLA. The PLA foci were counted from at least 20 cells. The data represent the mean ± SD from three independent experiments. The statistical significance is indicated by p-value (siControl vs sip85α, EGF stimulated). Related to Main Figure 4

**Figure S4.**
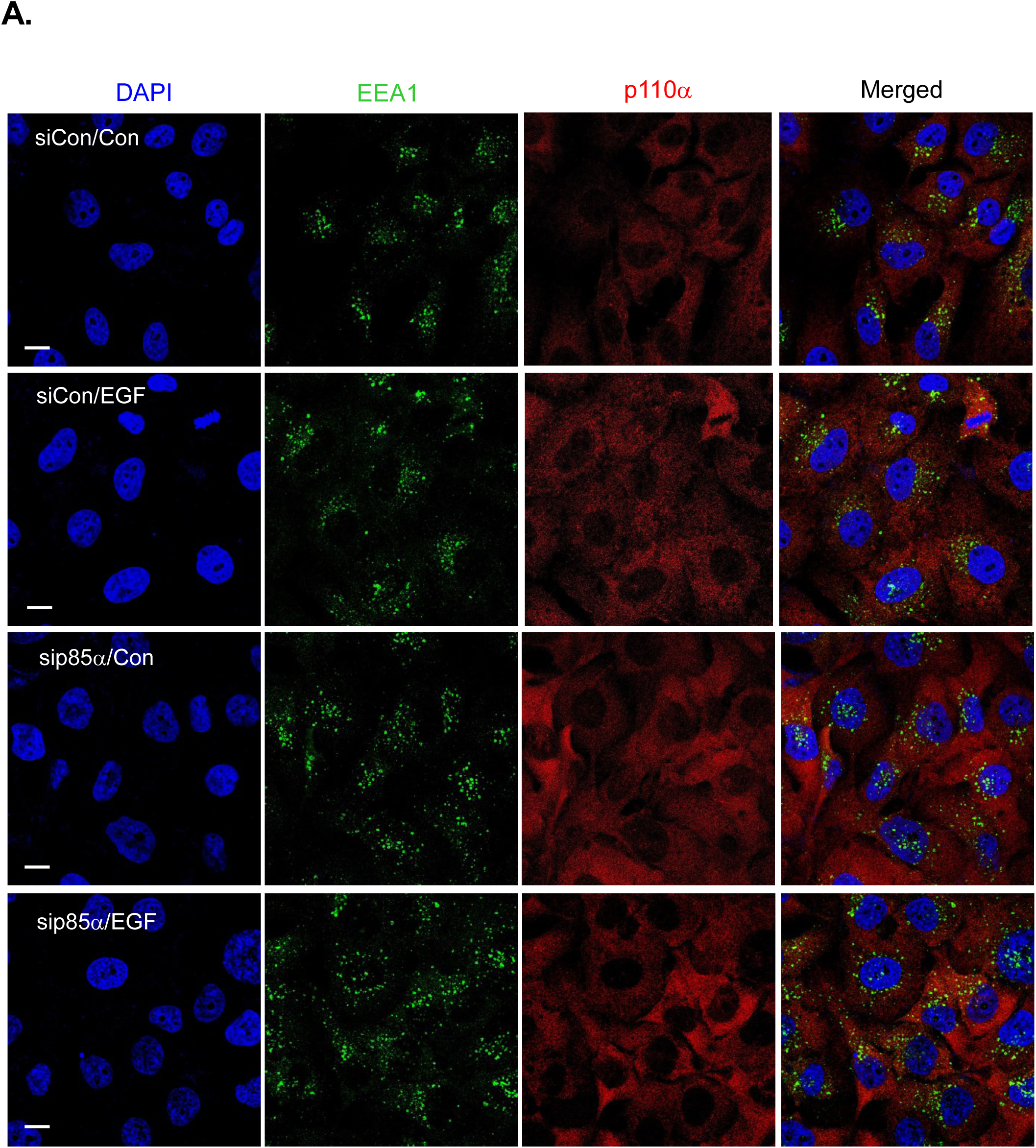
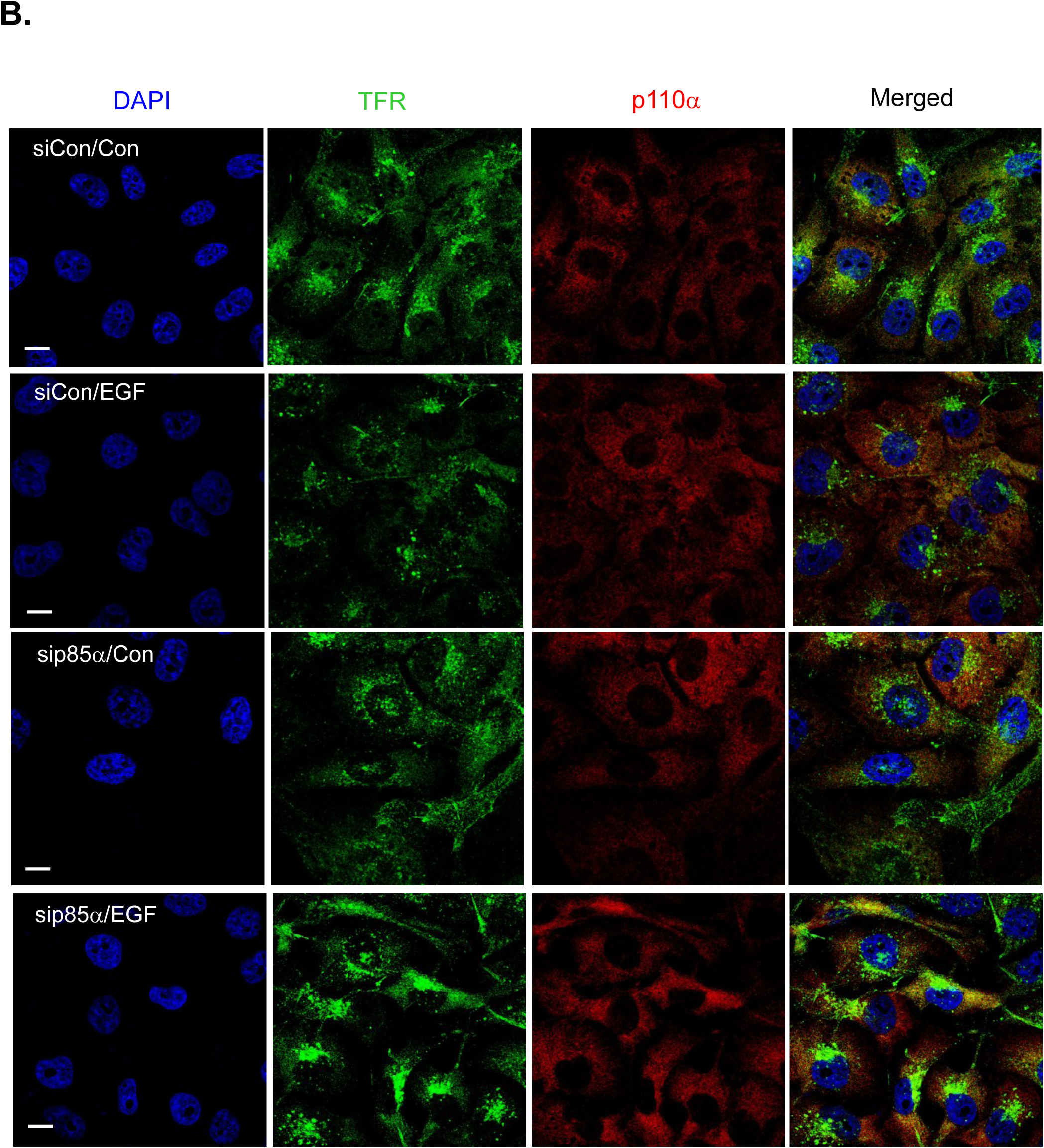
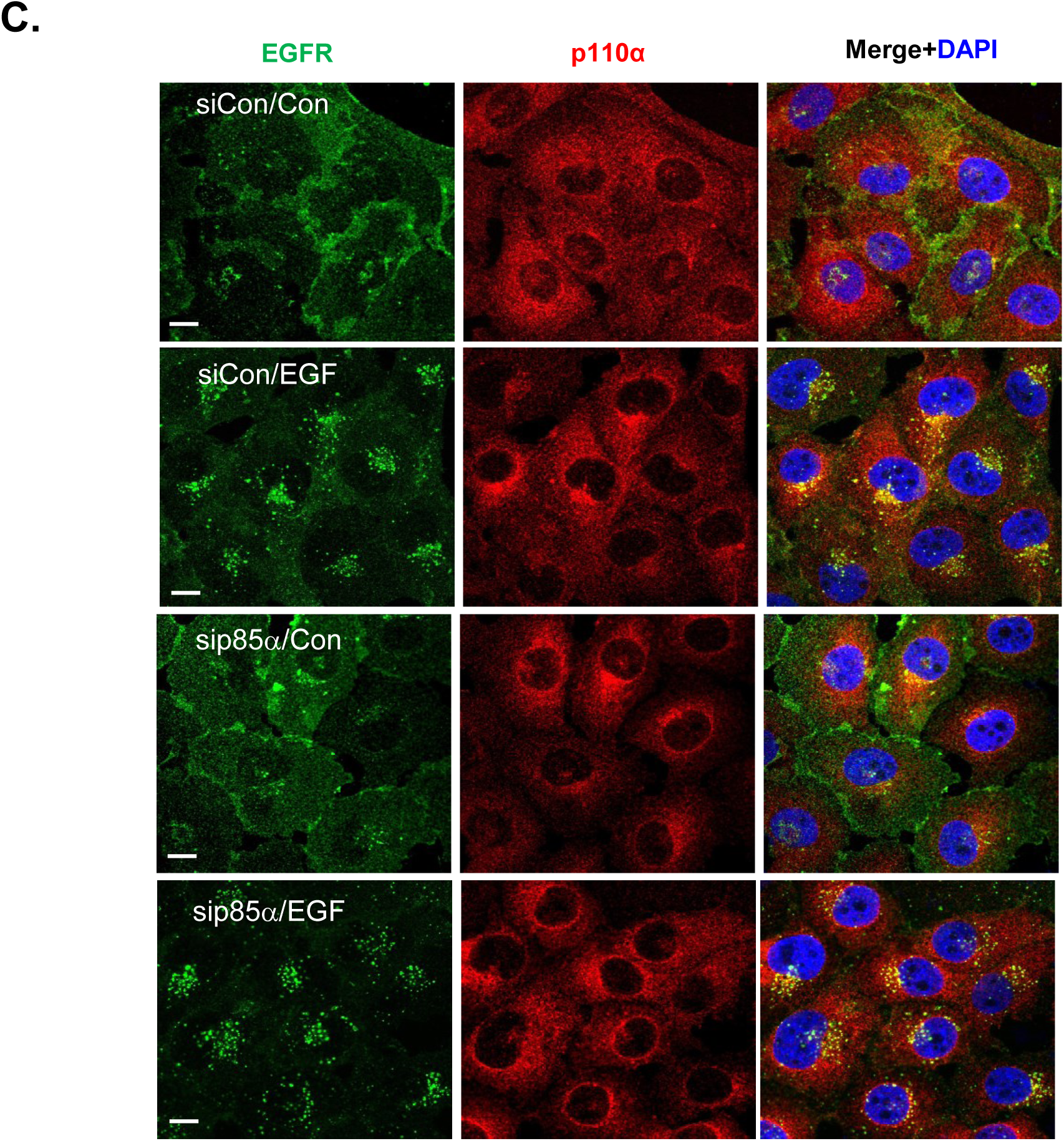
Enhanced p110α Co-localization with Endosomes and Association with EGF-stimulated Receptor Tyrosine Kinases upon p85α KD. **A, B, increased co-localization of p110α catalytic subunit with endosomes in response to EGF stimulation and p85α KD.** MDA-MB-231 cells were transfected with control siRNA or p85α siRNAs. 48-72 hours post-transfection, cells were stimulated with EGF before fixing the cells with 4% PFA. The images shown are representative images; part of these are shown in Figure 4 (B, C). **C, D, increased association of p110α catalytic subunit with EGF-stimulate EGFR upon p85α KD.** MDA-MB-231 cells were transfected with control siRNAs or p85α siRNAs. 48-72 hours post-transfection, cells were stimulated with EGF for 5 minutes before fixing the cells with 4% PFA for IF using antibodies specific for p110α (red) and EGFR (green). The images shown are representative images; part of these are shown in Figure 4 (E, F). Related to Main Figure 4

**Figure S5.**
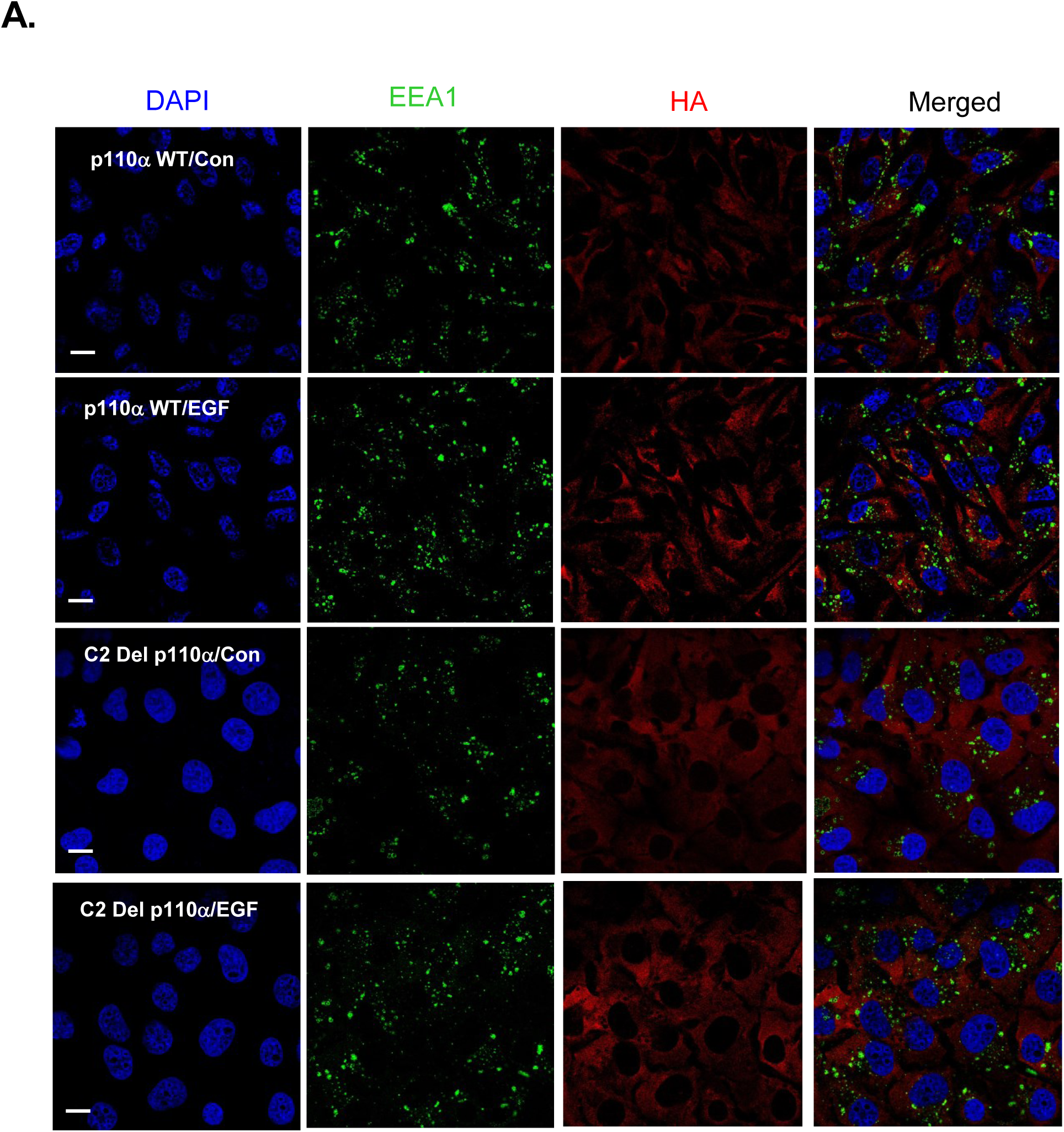
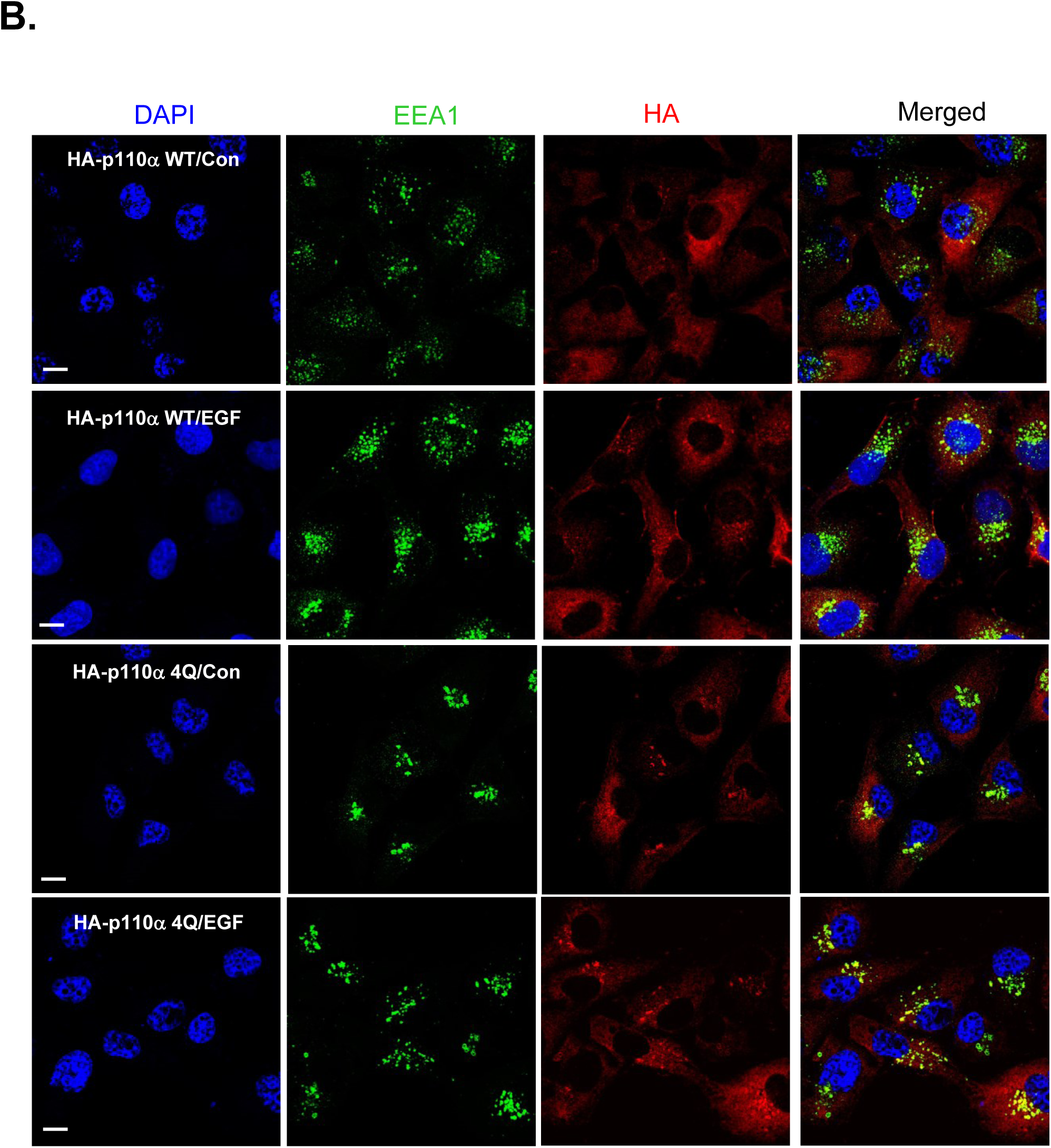
The p110α C2 domain-PI3P Interaction is Required for p110α Integration into Endosomes. **A,** MDA-MB-231 cells stably expressing HA-p110α WT or C2 domain deletion mutant HA-p110α were stimulated with EGF for 5 minutes before fixing the cells with 4% PFA. The cells were analyzed by IF with antibodies for HA (red) and EEA1 (green). The images shown are the representative images and are the part of Figure 5 (A). **B,** MDA-MB-231 cells stably expressing HA-p110α WT or HA-p110α 4Q mutant were stimulated with EGF for 5 minutes before fixing the cells with 4% PFA. The cells were analyzed by IF with antibodies for HA (red) and EEA1 (green). The images shown are representative images; part of these are shown in Figure 5 (F). Related to Main Figure 5

**Figure S6.**
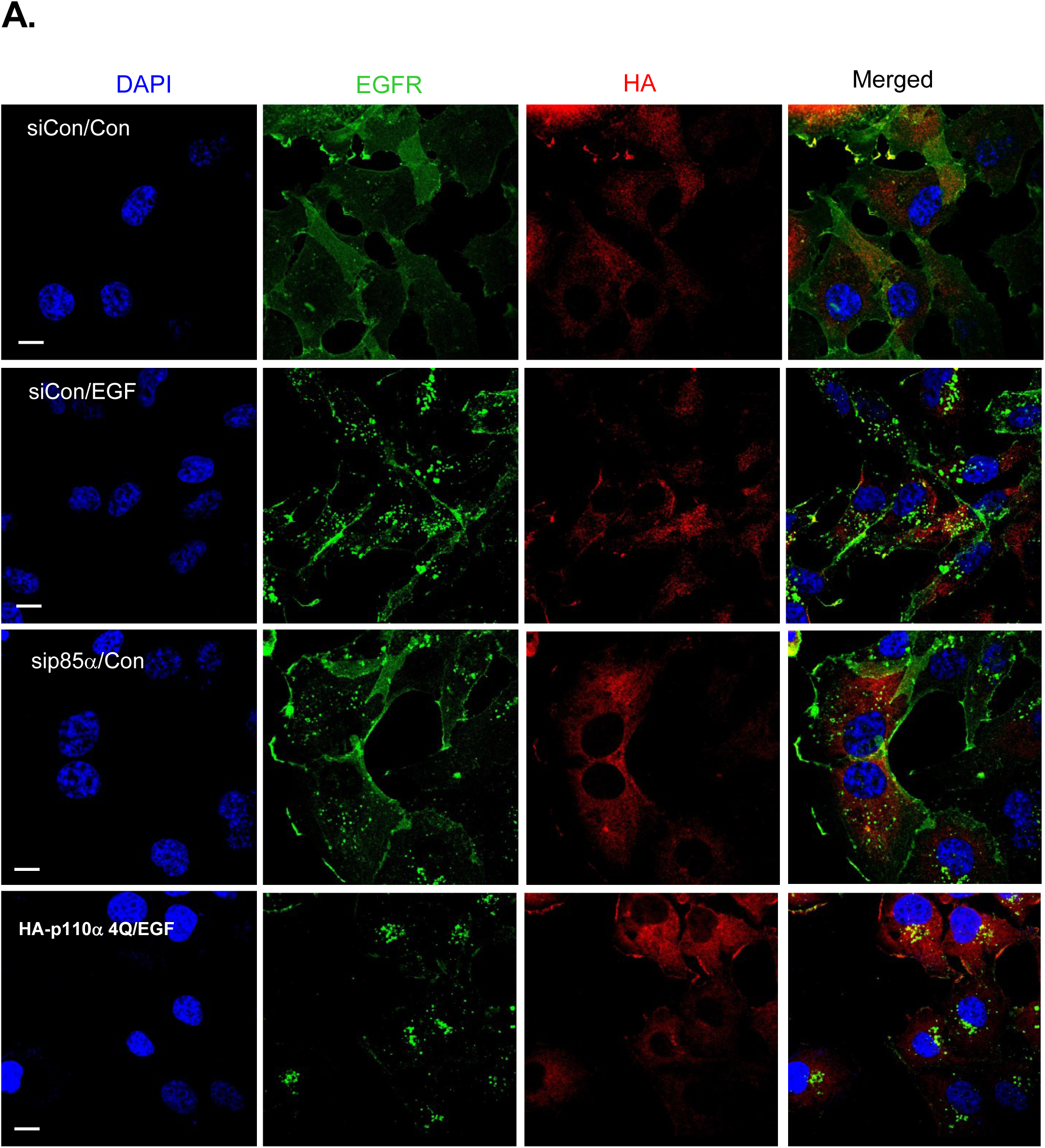
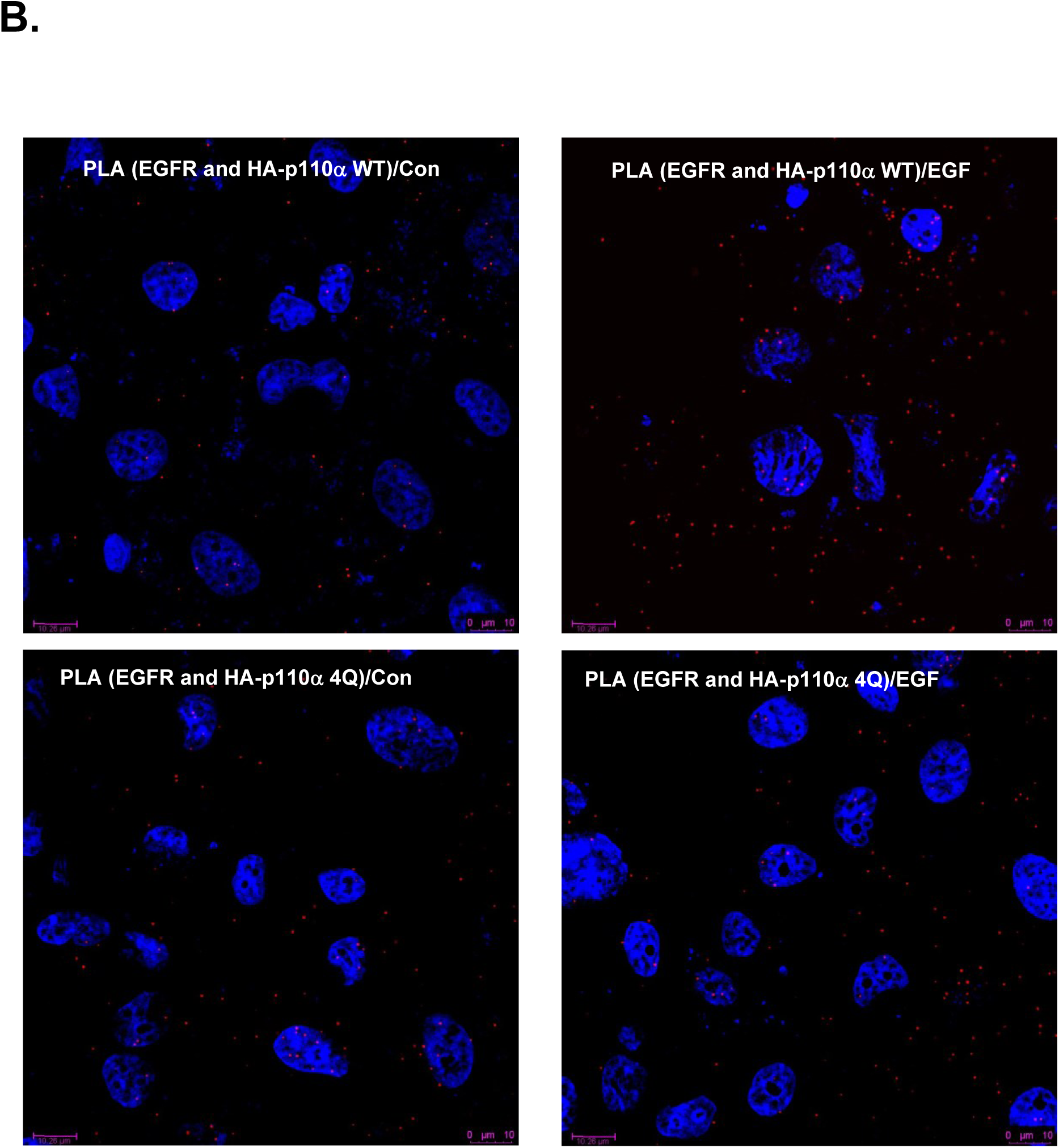
The p110α C2 domain-PI3P Interaction is Required for p110α Association with EGF-stimulated EGFR. **A,** MDA-MB-231 cells stably expressing HA-p110α WT or HA-p110α 4Q mutant were stimulated with EGF for 5 minutes before fixing the cells with 4% PFA. The cells were analyzed by IF with antibodies for HA (red) and EEA1 (green). The images shown are representative images; part of these are shown in Figure 5 (F). Related to Main Figure 6

## Notes

### Competing Interest Statement

The authors have declared no competing interest.

